# Spindle minus-end microtubule protrusions perturb force balance and delay chromosome segregation

**DOI:** 10.64898/2026.11.12.766267

**Authors:** Bojun Zhou, Zachary R. Gergely, Daniel G. Steckhahn, Taylor Mahoney, Lucas Swallow, J. Richard McIntosh, Mary Williard Elting, Meredith D. Betterton

## Abstract

Mitotic spindle function depends on multiple forces that are difficult to measure in cells. In fission yeast, the spindle assembles and elongates within an intact nuclear envelope. Without the kinesin-14/Pkl1, spindle microtubule minus ends detach from the spindle pole body and are pushed beyond it by kinesin-5/Cut7 sliding, forming protrusions that deform the nuclear envelope. We quantified and modeled protrusion dynamics. We found that as protrusions grew, the spindle often shortened, and the extent of shortening correlated with loss of spindle Cut7. Cells with protrusions showed a delay in anaphase onset. A force-balance model can explain fast and slow protrusion length changes, and inferred overlap length at force balance estimated the midzone sliding force density. After laser ablation, severed protrusions retracted and stopped near the cut, consistent with nuclear envelope-driven sliding. The results show the importance of minus-end anchoring for spindle mechanics and how protrusions can probe mechanical coupling between the spindle and nuclear envelope.

## Introduction

The mitotic spindle is a force-generating microtubule structure that organizes and segregates chromosomes prior to cell division ^1–5^. Spindle organization depends on microtubule polymerization and depolymerization, molecular motor activity, microtubule crosslinking and regulating proteins, chromosome attachments, and interactions with cell boundaries^3–14^. Motor proteins are typically required for spindle force generation and organization. In the force-balance model of spindle organization, kinesin-5 motors typically crosslink and slide antiparallel microtubules apart to generate outward forces that elongate the spindle, while kinesin-14 motors and dynein can generate inward forces that oppose spindle-pole separation^6,7,9–11,15–21^. This force balance helps determine spindle length and architecture, and perturbing it can disrupt spindle organization and compromise chromosome segregation^9,11,19,22^. However, the magnitude of individual contributions to spindle force remains difficult to determine in cells, because they act within an interconnected spindle ^2,23–25^.

The fission yeast *Schizosaccharomyces pombe* provides a useful system for studying spindle force balance. Its relatively simple bipolar spindle forms between two spindle pole bodies (SPBs), while the spindle assembles and elongates within an intact nuclear envelope ^26–28^. Sliding force to separate spindle poles is generated by the kinesin-5 Cut7 and kinesin-6 Klp9, opposed by the kinesin-14 motors Pkl1 and Klp2^18,29–34^. Remarkably, while Cut7 is considered essential for initial bipolar spindle assembly, loss of Cut7 can be rescued if Pkl1 is also deleted ^18,20,31,33,35–37^. In addition to its role in spindle force balance, Pkl1 forms a complex with Msd1 and Wdr8 to anchor spindle-microtubule minus ends at the SPBs ^37,38^. Loss of any member of the Pkl1-Msd1-Wdr8 complex disrupts normal spindle-pole organization and allows spindle microtubules to detach from and extend beyond the SPBs^37,38^. Spindle microtubule minus-end protrusions form in *pkl1* Δ cells, driven by Cut7-dependent sliding of unanchored spindle microtubules ^37,39^. Thus, when spindle microtubule minus-end anchoring is disrupted, a subset of microtubules can have their minus-ends driven beyond the spindle pole by Cut7-generated sliding.

These spindle microtubule minus-end protrusions provide a system to assess forces that are difficult to separate within an intact bipolar spindle. The fraction of cells showing protrusions has been used to determine changes in Cut7-generated sliding force. Truncations of the Cut7 C-terminal tail and mutations of its phosphorylation sites appear to lower sliding force generation, because temperature-sensitive spindle assembly, spindle length, and protrusion frequency decrease in these mutants ^40–42^. Because protrusion microtubules extend within the intact nucleus and deform the nuclear envelope as they elongate^18,37–39,42^, the dynamics of protrusions reflects both spindle-generated and nuclear-envelope forces. Previous work has further shown that the nuclear envelope can mechanically influence spindle shape and elongation^13,28,43–47^. Complementary laser-ablation studies have used acute mechanical perturbation to reveal forces that are otherwise balanced within the intact spindle. Severing the *S. pombe* spindle midzone causes the spindle poles to move inward and disrupts normal spindle elongation^21,48,49^. Protrusion formation and laser ablation therefore provide ways to assess mitotic force balance that are otherwise difficult to resolve.

Previous studies established that protrusions require Cut7 sliding ^39,41,42^. A mathematical force-balance model in which Cut7-driven outward sliding is opposed by the force of nuclear-envelope deformation qualitatively resembled experimentally measured protrusion length dynamics^42^. However, protrusion dynamics have not been systematically measured. Shortening of the spindle as a protrusion formed was observed^41,42^, but it remains unclear how protrusion dynamics affect spindle force balance, whether protrusions alter mitotic timing, or how protrusions respond to perturbation. Here, we combine quantitative live-cell imaging, quantitative image analysis, laser ablation, and computational force-balance modeling to characterize effects of protrusions on spindle force balance and mitotic progression. Protrusion growth and retraction showed fast and slow speeds, and protrusion extension was often associated with reversible spindle shortening and Cut7 redistribution. Cells with protrusions entered anaphase later than cells without protrusions. We also directly perturb protrusion force balance by severing protruding microtubules and show that mathematical modeling of protrusions can be used to estimate the forces generated by spindle motors. The results suggest that protrusions can serve as a probe of Cut7 force generation and spindle-nuclear envelope mechanical coupling.

## Results

### Spindle minus-end microtubule protrusions do not account for the reduced spindle length and elongation speed caused by Pkl1 loss

In a normal bipolar spindle in fission yeast, microtubule minus ends remain associated with the spindle pole bodies (SPBs). Spindle microtubule minus-end protrusions are formed when microtubules detach from the spindle pole. Protrusions occur in the absence of Pkl1, a kinesin-14 motor that helps anchor spindle microtubule minus ends at the SPBs and opposes the outward force generated by Cut7^18,37,39^. Protrusions occur in nearly 80% of mitotic *pkl1* Δ cells ^39,41,42^. To visualize protrusion dynamics, we performed live-cell confocal fluorescence imaging of *pkl1+* and *pkl1* Δ cells expressing the SPB marker Sid4-mCherry and the kinesin-5 motor Cut7-GFP (Figure 1A, B). In wild-type cells, Cut7-GFP localized along the spindle, including in the interpolar spindle and strongly near the SPBs, producing a characteristic pattern of two bright spots connected by dimmer midzone fluorescence (Figure 1A, C). In *pkl1* Δ cells with protrusions, additional Cut7-GFP was visible near the protrusion tip (Figure 1B, D). Consistent with the minus-end-directed activity of Cut7^40,50–52^, Cut7-GFP marks the microtubule minus end ^37,39,42^ to track the protrusion position relative to the Sid4-mCherry labeled SPB.

**Figure 1.**
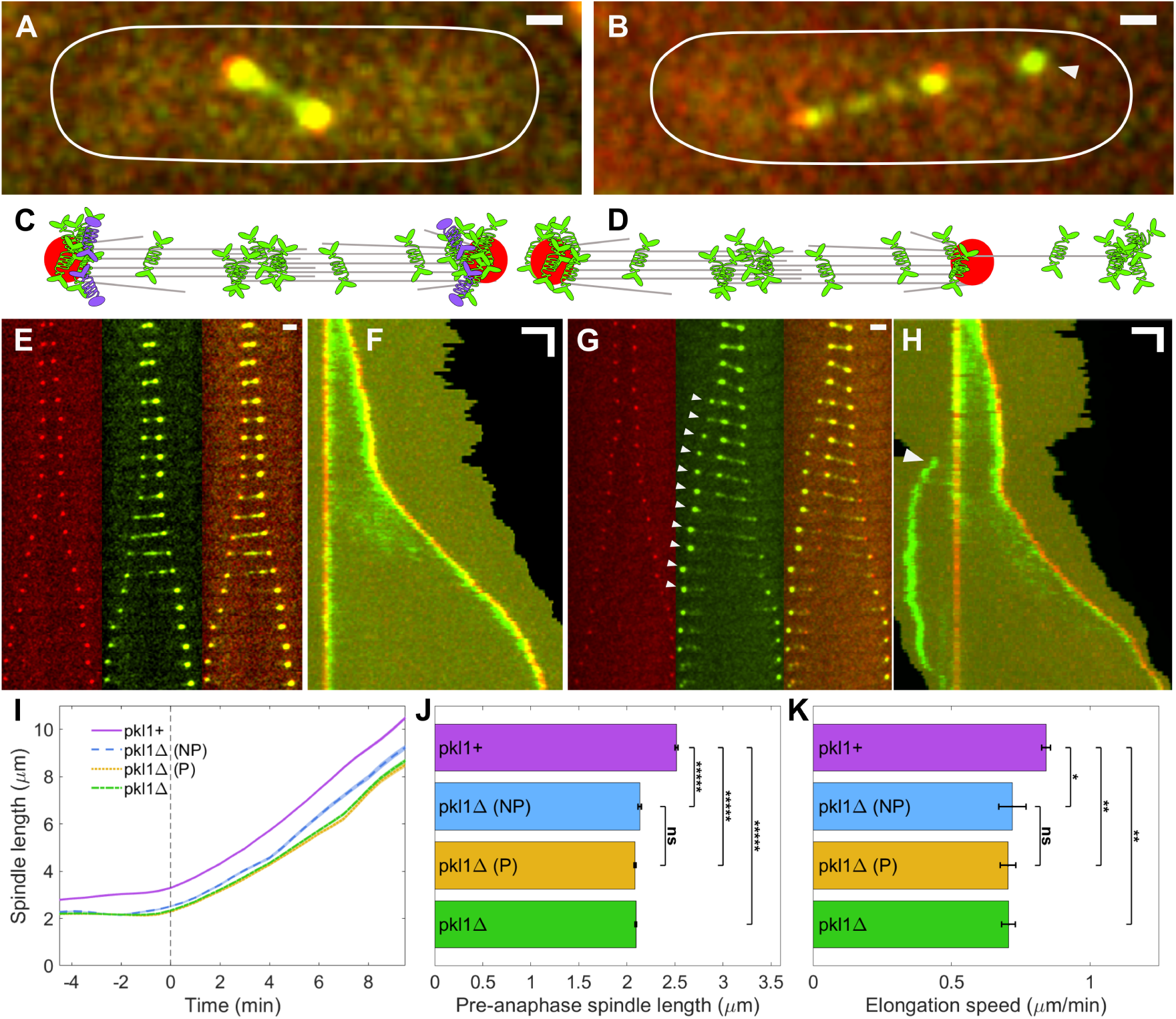
Spindle microtubule minus-end protrusions and spindle dynamics in *pkl1+* and *pkl1* Δ cells. **(A, B)** Confocal fluorescence micrographs of mitotic *pkl1+* **(A)** and *pkl1* Δ **(B)** cells showing SPBs (red, Sid4-mCherry) and Cut7-GFP (green), shown as maximum-intensity projections of confocal stacks. Cell boundaries are outlined by solid white lines, and the protrusion tip in B is indicated by white arrowheads. Scale bars, 1 *µ*m. **(C, D)** Schematics of a *pkl1+* spindle **(C)** showing Cut7 (green) and Pkl1 (purple) motors, microtubules (gray), and SPBs (red), and a *pkl1* Δ spindle **(D)** with a microtubule minus-end protrusion extending past the SPB with tip-localized Cut7. **(E)** Image sequence of a mitotic *pkl1+* cell showing SPBs (red, Sid4-mCherry, left), Cut7-GFP (green, center), and merge (right), shown as maximum-intensity projections of a confocal stack. Time between images, 72 s; scale bar, 2 *µ*m. **(F)** Two-color kymograph of the cell shown in E, constructed with one SPB fixed, creating a stationary red vertical line on the left. Dark regions of zero fluorescence intensity (upper right of image) indicate areas where the axis for kymograph projection extends beyond the cell segmentation boundaries (Methods). Scale bars, 2 *µ*m and 2 min. **(G)** Image sequence of a mitotic *pkl1* Δ cell, as in E. White arrowheads indicate protrusion tips with localized Cut7-GFP. Time between images, 70 s; scale bar, 2 *µ*m. **(H)** Two-color kymograph of the cell shown in G, constructed as in F. The white arrowhead marks the protrusion. Scale bars, 2 *µ*m and 2 min. **(I–K)** Quantification of spindle dynamics showing average spindle length as a function of time **(I)**, where the vertical dashed line denotes the time of anaphase B onset (*t* = 0 min); mean pre-anaphase spindle length **(J)**, with individual data points representing single frames sampled prior to anaphase B onset; and mean anaphase B spindle elongation speed **(K)** for *pkl1+* (*N* = 29), *pkl1* Δ-NP (*N* = 10), *pkl1* Δ-P (*N* = 32), and *pkl1* Δ (*N* = 42) cells. Error bars in J and K are standard error of the mean (SEM). Statistical significance was determined using one-way ANOVA followed by post-hoc Fisher’s LSD pairwise comparisons; \*\*\*\*\**p <* 0.00001, \*\**p <* 0.01, \**p <* 0.05, ns = not significant.

To measure spindle and protrusion dynamics, we constructed two-color kymographs using the Fission-Yeast Spindle Kymograph Analyzer (FYSKA) (Figure 1E–H and Videos 1–4) ^53^. In wild-type cells, Cut7-GFP colocalized with the SPBs throughout mitosis (Figure 1E, F). In early anaphase B, Cut7 was more visible on the interpolar spindle (Figure 1E, F), consistent with previous work ^40,42^. In *pkl1* Δ cells, Cut7 localization before protrusion formation was similar to that in wild-type cells. Once a protrusion formed, Cut7 accumulated at the protrusion tip (Figure 1G, H). Over time, Cut7 signal at the SPB near the protrusion decreased (Figure 1G, H), as observed previously ^42^. Thus, protrusion formation was accompanied by Cut7 redistribution toward the protrusion tip.

Protrusion microtubules detach from the SPBs and deform the nuclear envelope, which might alter spindle force balance. Therefore, we asked whether the presence of a protrusion was associated with changes in spindle dynamics. We quantified pre-anaphase spindle length and anaphase B elongation speed in *pkl1+* and *pkl1* Δ cells, and analyzed in the mutant population both cells without (*pkl1* Δ-NP) and with visible protrusions (*pkl1* Δ-P) (Figure 1I–K). When we compared the average spindle length over time, both *pkl1* Δ populations had shorter pre-anaphase spindles and slower subsequent elongation than wild-type (Figure 1I-K and Figure S1A–D). However, the presence or absence of a protrusion did not give a statistically significant difference in either spindle length or elongation speed (Figure 1J, K). Thus, the reductions in average spindle length and elongation speed are associated with loss of Pkl1 as previously observed^9,32^ and not the protrusion. This decrease in spindle length is counterintuitive, because removing a minus-end-directed motor would be expected to lengthen the spindle. Pkl1 may instead help maintain spindle length by aligning spindle microtubules so that Cut7 exerts outward force more effectively, as proposed for kinesin-14 in budding yeast ^54^. Pkl1 may also regulate microtubule nucleation ^35,36^, which could change the number of spindle microtubules available for sliding

### Spindle protrusions deform the nuclear envelope and are associated with delayed anaphase onset

To investigate how microtubule protrusions alter nuclear envelope (NE) shape, we imaged cells with both microtubule (mCherry-Atb2) and NE (Cut11-GFP) markers (Figure 2A). Because protruding microtubules extend past the SPB, they deform the NE ^37–39^. The distorted NE generates an inward force that opposes outward motor-driven sliding forces, altering the spindle force balance (Figure 2B). Because protrusions often form pre-anaphase, we asked whether protrusions impacted spindle positioning or altered the timing of anaphase onset.

**Figure 2.**
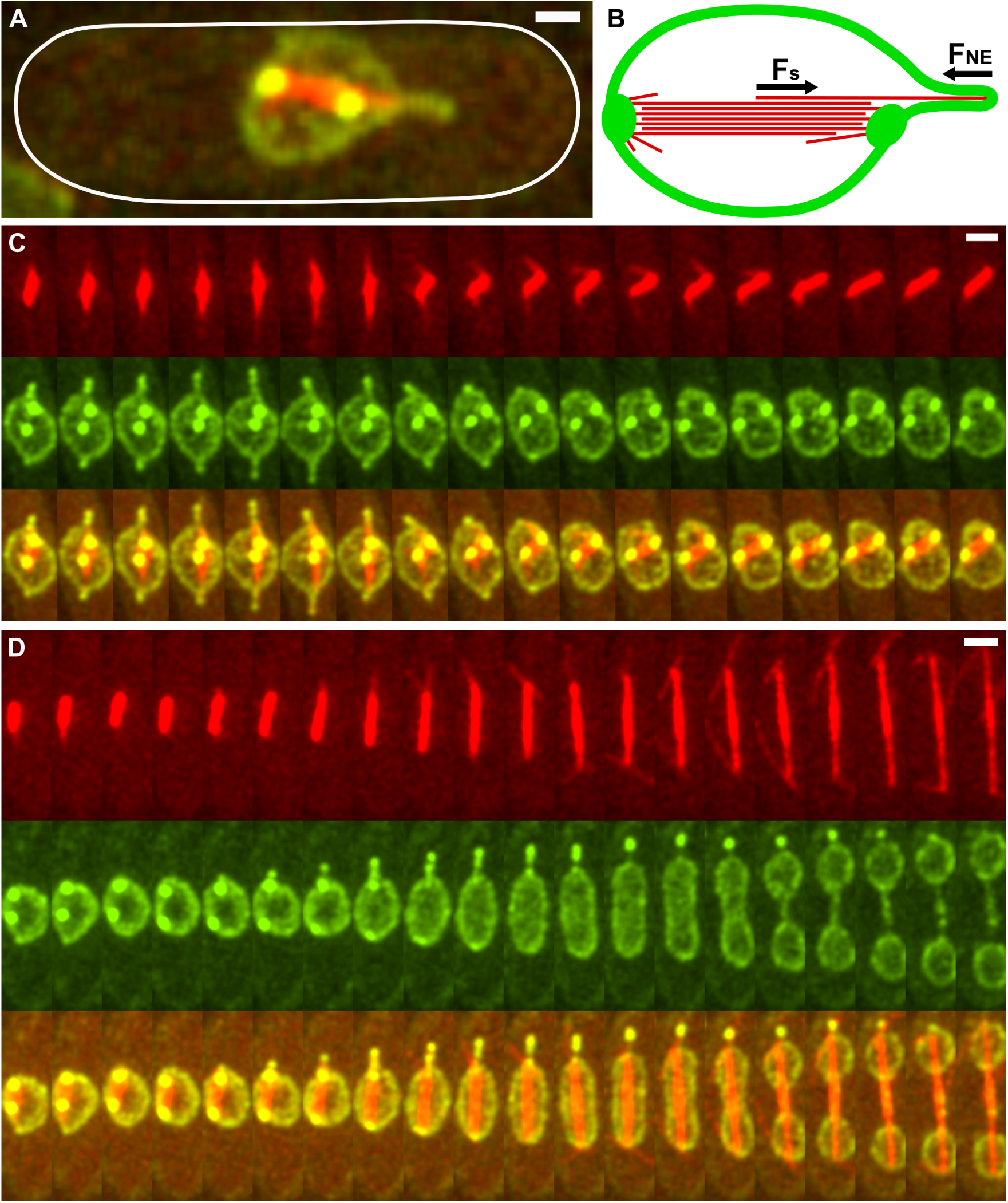
Spindle microtubule minus-end protrusions deform the nuclear envelope. **(A)** Fluorescence image of a spindle with a protrusion that deforms the nuclear envelope showing spindle microtubules (red, mCherry-Atb2) and the NE and SPBs (green, Cut11-GFP), shown as a maximum-intensity projection of a confocal stack. The cell boundary is outlined by solid white lines. Scale bar, 1 *µ*m. **(B)** Schematic of spindle microtubules with a protrusion (red) SPBs (green ovals), NE (green), and forces on the protrusion microtubule. **(C, D)** Image sequences of *pkl1* Δ cells showing spindle microtubules (red, mCherry-Atb2, top), NE and SPBs (green, Cut11-GFP, center), and merge (lower), shown as maximum-intensity projections of a confocal stack. The pre-anaphase cell **(C)** shows two protrusions that extend and deform the nuclear envelope, while the spindle axis rotates. During anaphase spindle elongation **(D)**, the NE deforms around the protrusion microtubules as its shape changes. Time between images, 50 s; scale bars, 2 *µ*m.

Protrusions induced dynamic, often asymmetric NE deformations throughout early mitosis (Figure 2C, D). In some cells, the NE distorted asymmetrically around a protrusion, forming a tighter interface on one side than the other (Figure 2C, top protrusion). This cell showed rotation of the spindle within the nucleus and the protrusion at an angle from the spindle (Figure 2C). This may reflect an unbalanced lateral force from the NE. In another cell, protrusions persisted as the cell entered anaphase (Figure 2D). We observed that the NE remained distorted around the protruding microtubules even as the main nuclear body underwent its typical progression to a peanut and dumbbell shape during spindle elongation (Figure 2D).

To determine whether protrusions affected mitotic progression, we measured the time from mitotic entry to anaphase onset (Methods, Figure 3). Mitotic entry was defined by the initial accumulation of Cut7-GFP at the SPBs (Figure 3A), left white arrowhead. Anaphase A onset was measured by the initial separation of sister kinetochore Mis12-RFP signals toward the two SPBs (Figure 3A), middle white arrowhead. Anaphase B onset was determined by the transition from steady pre-anaphase spindle length to rapid elongation (Figure 3A), right white arrowhead ^40^. Cells lacking visible protrusions (*pkl1* Δ-NP) had both anaphase A and anaphase B onset timing similar to wild-type cells (*pkl1+*) (Figure 3B, C). In contrast, cells with protrusions (*pkl1* Δ-P) showed a 3-minute delay in entering both anaphase A and B (Figure 3B, C). We additionally computed the interval between anaphase A and anaphase B onset and found that it was similar across all cell groups (Figure 3D). Thus, in *pkl1* Δ cells, the presence of a visible protrusion was associated with a longer time from the start of spindle assembly to the start of chromosome segregation.

**Figure 3.**
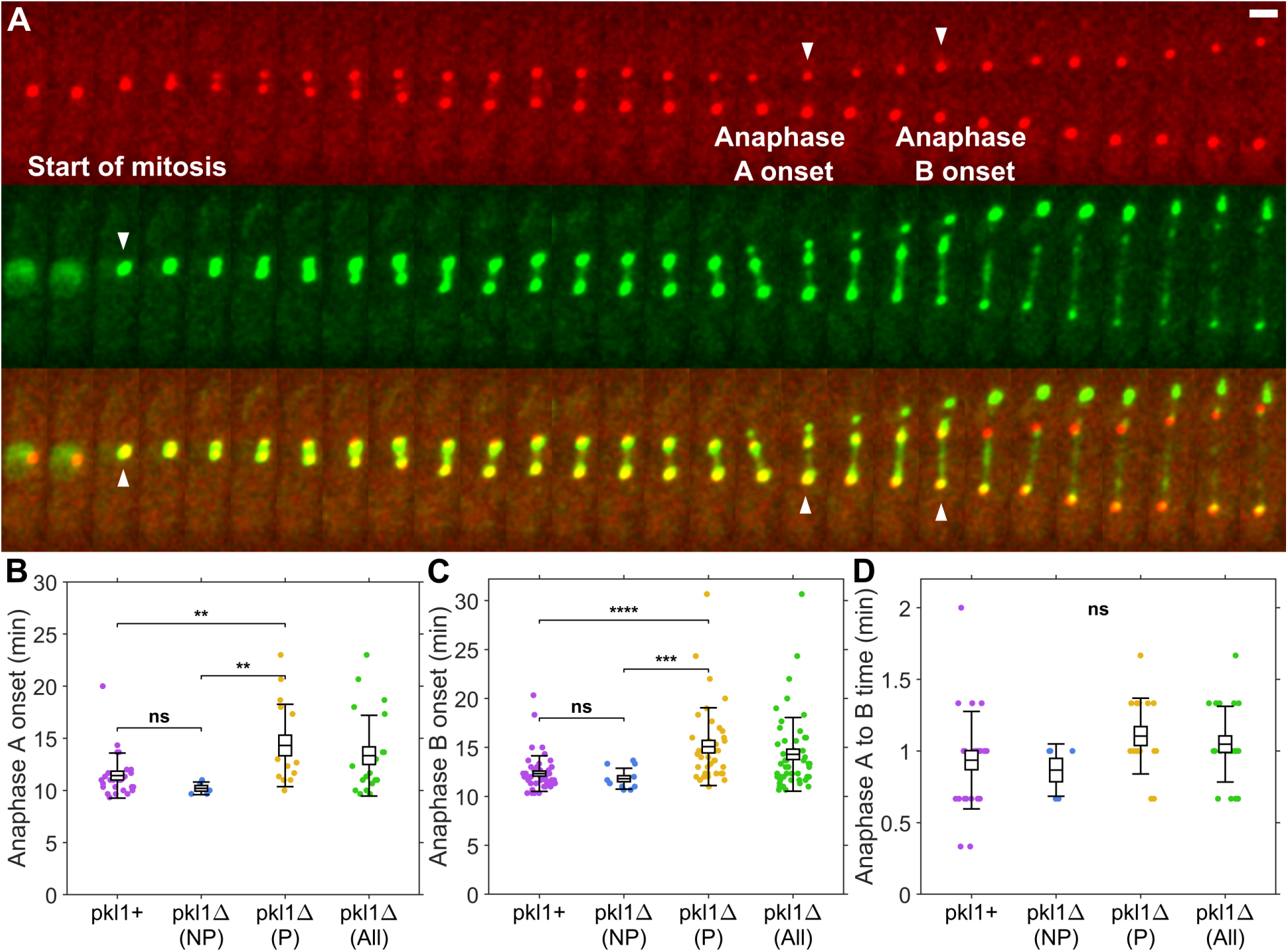
Spindle microtubule minus-end protrusions are associated with delayed anaphase onset. **(A)** Image sequence of a *pkl1* Δ cell showing SPBs and kinetochores (red, Sid4-mCherry and Mis12-RFP, top), Cut7-GFP (green, center), and merge (lower). Left, the start of mitosis was determined by the time when Cut7-GFP fluorescence was first visible at the SPBs (left white arrowhead). Anaphase A onset was the time when Mis12-RFP split toward the two SPBs (middle white arrowhead). Anaphase B onset was the time when more rapid SPB separation began (right white arrowhead). Time between images, 40 s; scale bar, 2 *µ*m. **(B)** Time from the start of mitosis to anaphase A onset for *pkl1+* (*N* = 26), *pkl1* Δ-NP (*N* = 5), *pkl1* Δ-P (*N* = 16), and *pkl1* Δ (*N* = 21) cells. **(C)** Time from the start of mitosis to anaphase B onset for *pkl1+* (*N* = 50), *pkl1* Δ-NP (*N* = 12), *pkl1* Δ-P (*N* = 38), and *pkl1* Δ (*N* = 50) cells. **(D)** Time from the onset of anaphase A to anaphase B for *pkl1+* (*N* = 26), *pkl1* Δ-NP (*N* = 5), *pkl1* Δ-P (*N* = 16), and *pkl1* Δ (*N* = 21) cells. Boxes show the mean and SEM; error bars show the standard deviation. Statistical significance was determined using one-way ANOVA followed by post-hoc Fisher’s LSD pairwise comparisons; \*\*\*\**p <* 0.0001, \*\*\**p <* 0.001, \*\**p <* 0.01, ns = not significant.

### Spindle length shortens during protrusion formation and recovers upon retraction, coinciding with a reduction in spindle Cut7 intensity

To investigate the interaction between spindle dynamics, protrusion formation, and NE shape, we first analyzed pre-anaphase spindles. Protrusion events occurred most frequently for short spindles when the NE is approximately spherical and confined the spindle prior to anaphase NE area expansion ^43,55^. To quantify spindle and protrusion dynamics, we tracked both SPBs using FYSKA^53^ and manually labeled the Cut7-GFP fluorescence associated with protrusion tips in every imaging frame where they were visible (Methods). In these three-spot images, we labeled the SPB adjacent to the protrusion the middle SPB and the SPB on the opposite side of the spindle the far SPB.

At the onset of protrusion formation, as the protrusion tip moved away from the middle SPB, the distance between the far SPB and middle SPB often decreased. Thus the bipolar spindle shortened as the protrusion extended (Figure 4A–G). Typically, the maximum protrusion length was simultaneous with the minimum spindle length. We noticed that protrusions sometimes rapidly shortened, which we denote fast retraction of the protrusion. During fast retraction, we often observed that spindle length increased (Figure 4A–G). These results suggest that protrusion elongation could alter spindle force balance reversibly.

**Figure 4.**
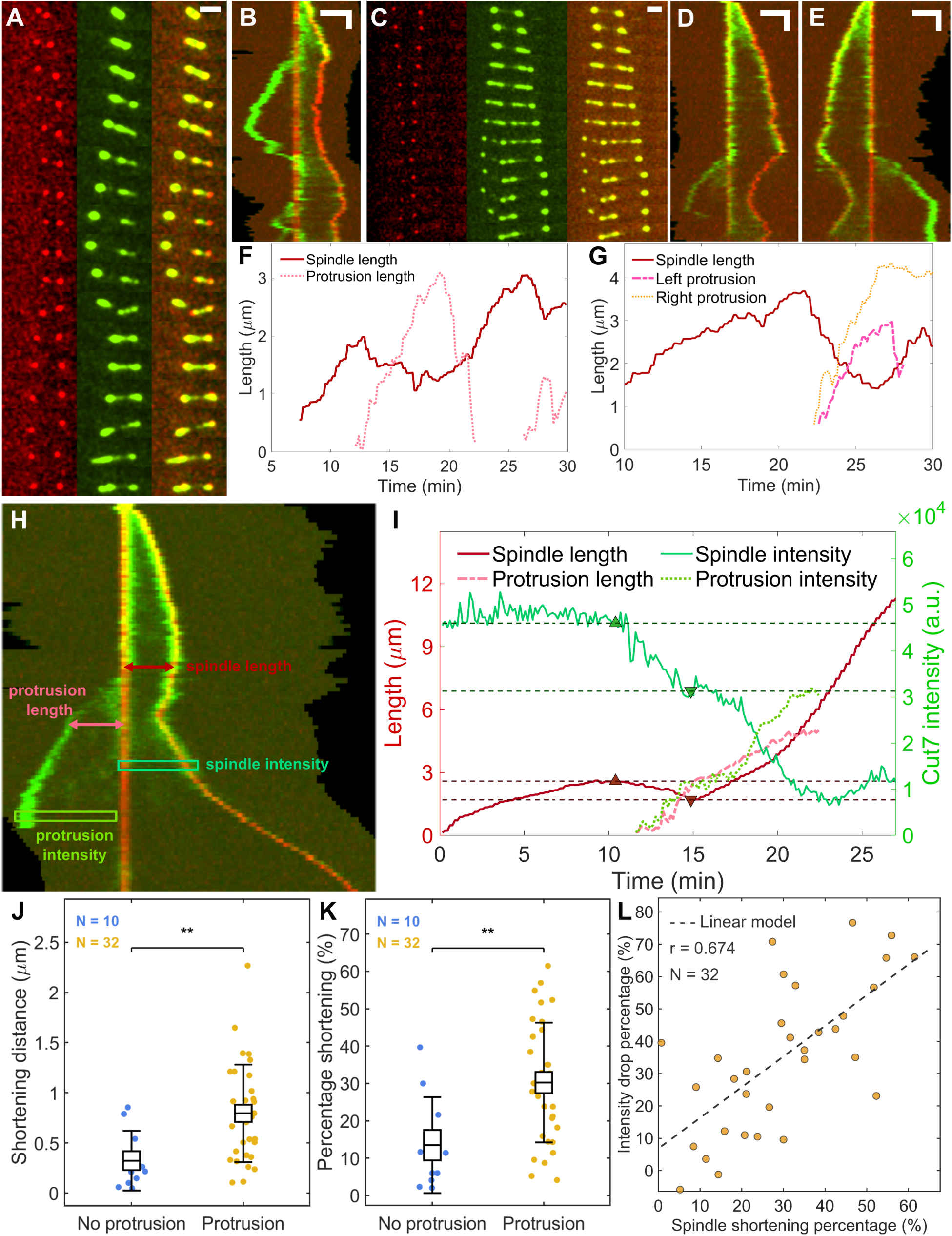
Cut7 relocalization to protrusion microtubules is correlated with spindle shortening during protrusion extension. **(A)** Image sequence of a *pkl1* Δ cell showing SPBs (red, Sid4-mCherry, left), Cut7-GFP (green, center), and merge (right), shown as maximum-intensity projections of a confocal stack. A protrusion extends from the left side of the spindle. Time between images, 75 s; scale bar, 2 *µ*m. **(B)** Kymograph of the cell shown in A. Scale bars, 2 *µ*m and 2 min. **(C)** Image sequence of a *pkl1* Δ cell, as in A. Two protrusions are visible on either side of the spindle. Time between images, 36 s; scale bar, 2 *µ*m. **(D, E)** Kymographs of the cell shown in C, with projection axes oriented to highlight the left protrusion **(D)** and the right protrusion **(E)**. Scale bars, 2 *µ*m and 2 min. **(F)** Spindle and protrusion length as a function of time for the cell shown in A and B. **(G)** Spindle and protrusion length as a function of time for the cell shown in C–E. **(H)** Kymograph of a mitotic *pkl1* Δ cell indicating measured spindle and protrusion length (dark and light red double-headed arrows) and boxes used to measure Cut7-GFP signal (teal-green and yellow-green). **(I)** Spindle and protrusion length (left axis, red) and Cut7-GFP intensity (right axis, green) as a function of time for the cell shown in H. Upward and downward triangles indicate local maxima and minima. Dashed lines indicate the spindle shortening distance and change in Cut7-GFP intensity. **(J, K)** Quantification of absolute **(J)** and relative **(K)** spindle shortening for cells without (*N* = 10) and with (*N* = 32) protrusions. Boxes show the mean and SEM; error bars show the standard deviation. Statistical significance was determined using one-way ANOVA; \*\**p <* 0.01. **(L)** Plot of relative spindle Cut7-GFP intensity drop as a function of relative spindle shortening (*N* = 32). The black dashed line shows the linear regression; Pearson correlation *r* = 0.674, *p <* 0.0001.

Our observations suggested that relocalization of the kinesin-5 Cut7 could contribute to spindle shortening. Directed motility of Cut7-GFP from the spindle midzone and SPBs toward the protrusion tips was observable in our kymographs, consistent with previous results^42^. We hypothesized that as Cut7-GFP accumulated at the protrusion tip, the active motors were depleted from the spindle midzone, reducing outward sliding force and contributing to spindle shortening. To test this, in each image we measured spindle length, protrusion length, spindle Cut7-GFP intensity (including both SPBs and the midzone, Methods), and protrusion Cut7-GFP (including the whole protrusion, Methods) (Figure 4H). We then quantified the maximum spindle length before protrusion formation and the minimum after protrusion elongation, and determined the change in spindle Cut7-GFP intensity during this time. We found that spindle Cut7-GFP intensity decreased during spindle shortening (Figure 4I). We note that the decrease in spindle Cut7-GFP fluorescence was often visible before the protrusion tip could be tracked. This may occur due to resolution limits in identifying short protrusions or a delay in Cut7-GFP localization to the protrusion tip.

To further test whether protrusion formation and Cut7 relocalization contribute to spindle shortening, we compared to *pkl1* Δ cells without protrusions by measuring spindle length maxima and subsequent minima pre-anaphase (Methods, Figure S1G–I). This method that looks for spindle length changes found an average 0.32 *±* 0.09 *µm* decrease in *pkl1* Δ-NP cells. In contrast, *pkl1* Δ cells with protrusions had a significantly greater mean shortening of 0.80 *±* 0.09 *µm* (Figure 4J). Because this shortening was transient, it did not change the average pre-anaphase length (Figure 1J). Normalizing this as a shortening percentage (shortening distance divided by local maximum length) showed that *pkl1* Δ-P cells had an average of 30% *±* 3% shortening, significantly higher than *pkl1* Δ-NP cells with an average of 14% *±* 4% (Figure 4K). To assess how well Cut7 relocalization could explain spindle shortening, we found the correlation between the percentage of spindle shortening and the percentage drop in spindle Cut7-GFP intensity. The correlation had Pearson’s *r* = 0.674 (Figure 4L). This suggests that midzone Cut7 depletion contributes to spindle shortening during protrusion extension. The inward force exerted by the NE could also contribute to spindle shortening.

### High-temporal resolution imaging reveals fast and slow protrusion length changes

We observed that protrusion dynamics changed before and after anaphase onset. Pre-anaphase protrusions were highly dynamic, with varying speed during both growth and retraction. Protrusion growth typically began with rapid growth when the tip quickly extended, followed by slower extension (Figure 5A). Conversely, protrusion retraction initially proceeded slowly or remained nearly stationary before rapidly retracting back to the SPB (Figure 5B). In contrast, when a protrusion persisted past anaphase onset, its dynamics were usually slower. Anaphase protrusions typically remained beyond the SPB for over 10 minutes before slowly merging with the SPB (Figure S2).

**Figure 5.**
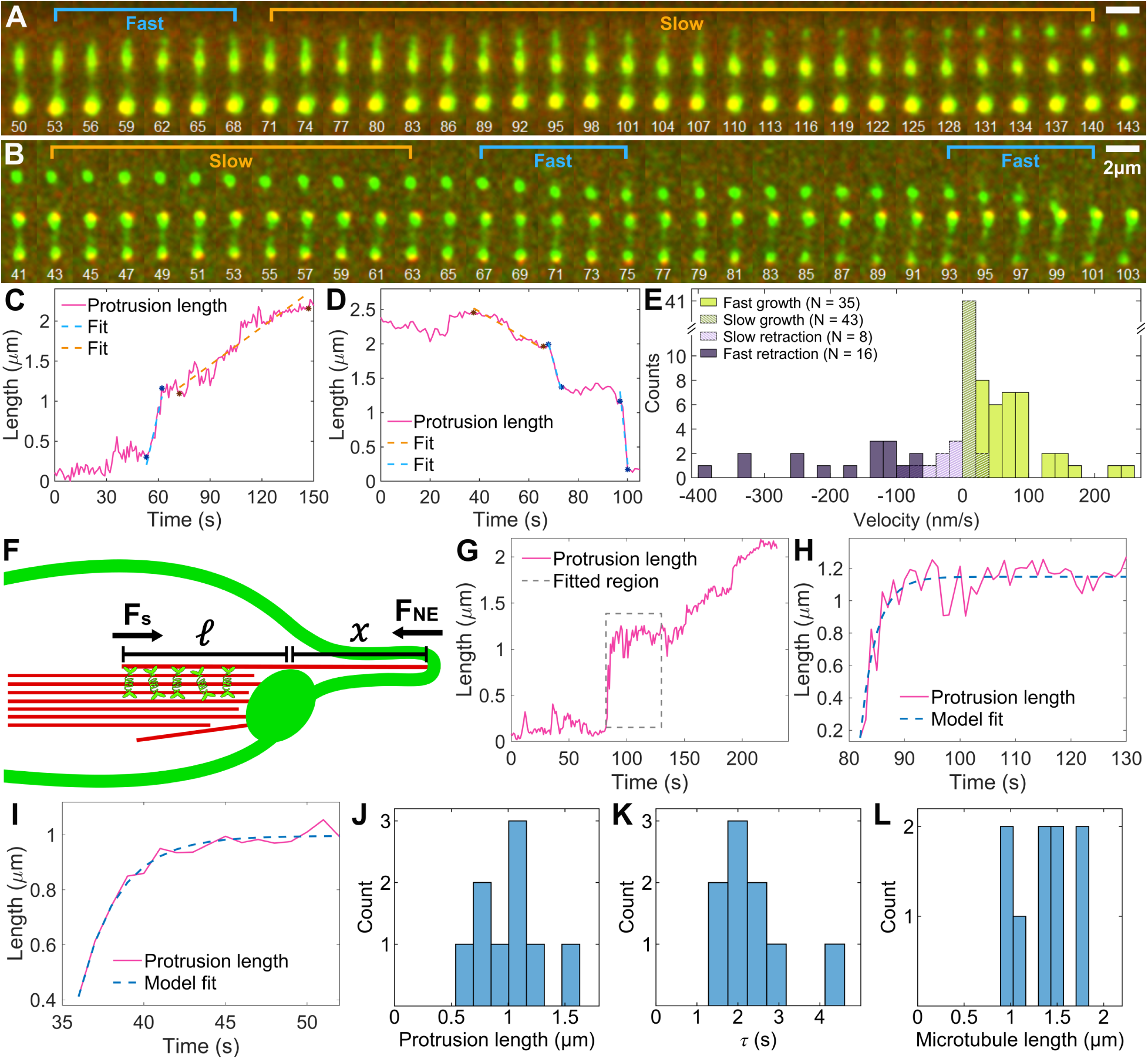
Pre-anaphase protrusions show fast and slow growth and retraction, a force-balance model can explain fast growth. **(A, B)** Fluorescence image sequences of *pkl1* Δ cells showing SPBs (red, Sid4-mCherry) and Cut7-GFP (green) during fast (light blue) and slow (orange) growth **(A)** and retraction **(B)**. Time shown in s. Images acquired every 1 s and shown every 3 s in A and 2 s in B; scale bar, 2 *µ*m. **(C, D)** Protrusion length as a function of time and linear fits of fast and slow protrusion growth **(C)** and protrusion retraction **(D)** for the cells shown in A and B. **(E)** Histogram of fitted protrusion speeds for fast growth (*N* = 35), slow growth (*N* = 43), slow retraction (*N* = 8), and fast retraction (*N* = 16) from cells showing growth (*N* = 28) and retraction (*N* = 15). **(F)** Schematic of the model of protrusion length dynamics. The protrusion microtubules experience force from the midzone *F*_s_ and nuclear envelope *F*_NE_. The midzone sliding force is proportional to the length of overlap region *ℓ*. **(G–H)** Example fit of the fast protrusion growth. Full trajectory (G) and fitted region (H). **(I)** Another example fit of the fast protrusion growth. **(J)** Histogram of force-balance protrusion length *x*_b_ from model fitting (*N* = 9). **(K)** Histogram of time constant *τ* from model fitting. **(L)** Histogram of total initial protrusion microtubule length *L* from model fitting.

We quantified protrusion growth and retraction velocities and categorized protrusion trajectories into four regimes: fast growth, slow growth, slow retraction, and fast retraction. We then fitted to determine the velocity of each event (Figure 5C, D, S3A–D) and determined the velocity distributions (Figure 5E, S3E–I). Slow growth and retraction had velocities 0 to 25 nm/s and *−*80 to 0 nm/s. Fast growth reached velocities up to *∼*250 nm/s, and fast retraction had the highest speeds, with velocities reaching *−*400 nm/s (Figure 5E, S3H). We compared these to measured mitotic microtubule polymerization dynamics in *S. pombe*. Typical microtubule growth speeds are 45-70 nm/s with events measured in the range 10-180, while typical shrinking speeds are 60-110 nm/s, with a range of 20-400^56–58^. The slow protrusion growth and retraction speeds fall in the typical dynamic instability range. The fast events are higher than the typical values, although fast retraction speeds did lie in the broader range of microtubule shrinking speeds that have been observed. The fast protrusion growth speeds were faster than has been observed for microtubule polymerization in fission yeast.

### A force-balance model of fast protrusion growth estimates midzone sliding force

To estimate the forces acting on protrusions, we extended a force-balance model of protrusion dynamics ^42^ (Methods). In this model, the total length of a protrusion microtubule is *L* = *x* + *ℓ*, where *x* is the length of the protrusion, the part of the microtubule that extends beyond the SPB, and *ℓ* is the length of the segment that overlaps the spindle midzone (Figure 5F). In this model, midzone motors push the protrusion outward with sliding force *F*_s_ = *f*_s_*ℓ*, proportional to overlap length, while the NE tube pushes the protrusion inward with a constant force *F*_NE_. We assume that fast length changes occur when the forces on the protrusion are unbalanced (the net force is nonzero). Assuming that microtubule polymerization kinetics are slow on this time scale, *L* is constant during fast events and any changes in protrusion length occur because sliding increases *x* and decreases *ℓ*. We neglect inward SPB motion during these events, because spindle shortening occurred over minutes and fast growth over seconds (Figure 4F, G). As the protrusion grows, the overlap and the sliding force decrease until the forces balance according to *f*_s_*ℓ*_b_ = *F*_NE_, which lets us find *x*_b_ and *ℓ*_b_ at mechanical balance. The model predicts that protrusion length approaches *x*_b_ exponentially with time constant *τ* = *γ/f*_s_, where *γ* is the protrusion drag coefficient (Methods). To determine how well this model describes fast protrusion growth, we fit the exponential solution to fast growth events, with fit parameters *x*_b_ and *τ* (Figure 5G–K, S4). The mean values were *τ* = 2.4 *±* 0.3 s and *x*_b_ = 1.0 *±* 0.1 µm, and these estimates along with low and high values determined from the standard deviation are shown in table 1.

**Table 1.** Fit results and estimated and derived parameters. Estimates are means. For *τ*, *x*_b_, *L*, *ℓ*_b_ (*N* = 9) and *v*_r_ (*N* = 13), low and high values are the mean minus and plus the standard deviation across events. For rows that depend on *κ*, low and high values are determined from the ends of the *κ* range, with *v*_r_ and *ℓ*_b_ at their estimates.

| Quantity | Low | Estimate | High |
|---|---|---|---|
| *Measured and fit* |  |  |  |
| Relaxation time *τ* (s) | 1.4 | 2.4 | 3.3 |
| Force-balance protrusion length *x*<sub>b</sub> (µm) | 0.7 | 1.0 | 1.3 |
| Retraction speed *v*<sub>r</sub> (nm/s) | 80 | 140 | 200 |
| *Derived, independent of κ* |  |  |  |
| Protrusion microtubule length *L* (µm) | 1.0 | 1.4 | 1.7 |
| Force-balance overlap length *ℓ*<sub>b</sub> (µm) | 0.20 | 0.33 | 0.46 |
| *Estimated and derived, dependent on κ* |  |  |  |
| Bending modulus *κ* (pN nm) | 80 | 230 | 660 |
| NE tube force *F*<sub>NE</sub> (pN) | 5.7 | 16.5 | 47 |
| Drag coefficient *γ* = *F*<sub>NE</sub>/*v*<sub>r</sub> (pN s/µm) | 41 | 118 | 340 |
| Sliding force density *f*<sub>s</sub> = *F*<sub>NE</sub>/*ℓ*<sub>b</sub> (pN/µm) | 17 | 50 | 140 |

To infer additional parameters, we also analyzed fast retraction events. We hypothesize that fast retraction occurs when the protrusion microtubule loses contact with the midzone, so *F*_s_ = 0. The retraction speed is then set by the NE force and drag alone, *v*_r_ = *F*_NE_*/γ*. The measured mean speed was *v*_r_ = 139 *±* 17 nm/s, calculated with 3 outliers with speed *v*_r_ *>* 300 nm*/*s excluded (Table 1, Figure S3H). Given the retraction speed, we can combine the force balance relation *ℓ*_b_*f*_s_ = *F*_NE_ with the fast retraction result *v*_r_ = *F*_NE_*/γ* to eliminate the NE force, finding *ℓ*_b_ = *v*_r_*τ*. As a result, the fits directly infer the force-balance overlap length *ℓ*_b_ and the protrusion microtubule length *L* = *x*_b_ + *ℓ*_b_. The mean values were *ℓ*_b_ = 0.33 *±* 0.05 µm, and *L* = 1.4 *±* 0.1 µm (Figure 5L, table 1). Protrusions typically began when the spindle length was about 3 m, giving an upper limit on the length of protrusion microtubules when the protrusion is first visible. All fits inferred microtubule lengths *L* below 3 µm. Thus, the inferred lengths fell within an independent limit set by spindle length at protrusion onset.

We can use these parameters inferred from the fits to estimate forces from midzone sliding. For the fission yeast NE, only the ratio of tension to bending modulus has been estimated ^59^. The NE bending modulus is thus uncertain, and we consider it to range from 80 to 660 pN nm, with an estimate 230 pN nm. We then can calculate the force from a nuclear envelope tube *F*_NE_ = 16.5 pN, with range 5.7 to 47 pN (Methods, Table 2). From force balance, we can then infer sliding force density *f*_s_ = *F*_NE_*/ℓ*_b_ *≈* 50 pN*/*µm, with range 17 to 140 pN/µm. For comparison, human Eg5 sliding apart a single antiparallel microtubule pair produced a plateau force that increased with overlap length at 4 pN/µm ^60^. Our estimate is 12 times this value, which may occur because the fission yeast midzone contains 3 to 5 antiparallel pairs, motors may crosslink between more than one pair and the protrusion, and the midzone motor density may be higher in the cell than in reconstitution experiments.

**Table 2.** Estimated NE tension and force for a range of bending modulus values.

| Basis | *κ* (pN nm) | *σ* (pN/nm) | *F*<sub>NE</sub> (pN) |
|---|---|---|---|
| Low, PC vesicles | 80 | 0.0052 | 5.7 |
| Estimate | 230 | 0.015 | 16.5 |
| High, ER and Golgi membranes | 660 | 0.043 | 47 |

### After laser ablation, protrusion microtubules move toward the SPB and stop near the ablation site

Our model suggests that protrusion length dynamics arise from a balance between outward sliding force and inward NE force. A protrusion fragment severed from the midzone has no overlap and therefore no sliding force, so the model predicts that it retracts at the fast retraction speed *v*_r_. To test this prediction, we used laser ablation to cut protrusion microtubules. To visualize how protrusion microtubules respond to ablation, we performed these experiments with two different fluorescent markers: a strain with Cut7-GFP to label the SPBs and protrusion tips and low-expression mCherry-Atb2 to label microtubules (Figure 6A), and a strain with GFP-Atb2 to label spindle microtubule and protrusion microtubules (Figure 6B).

**Figure 6.**
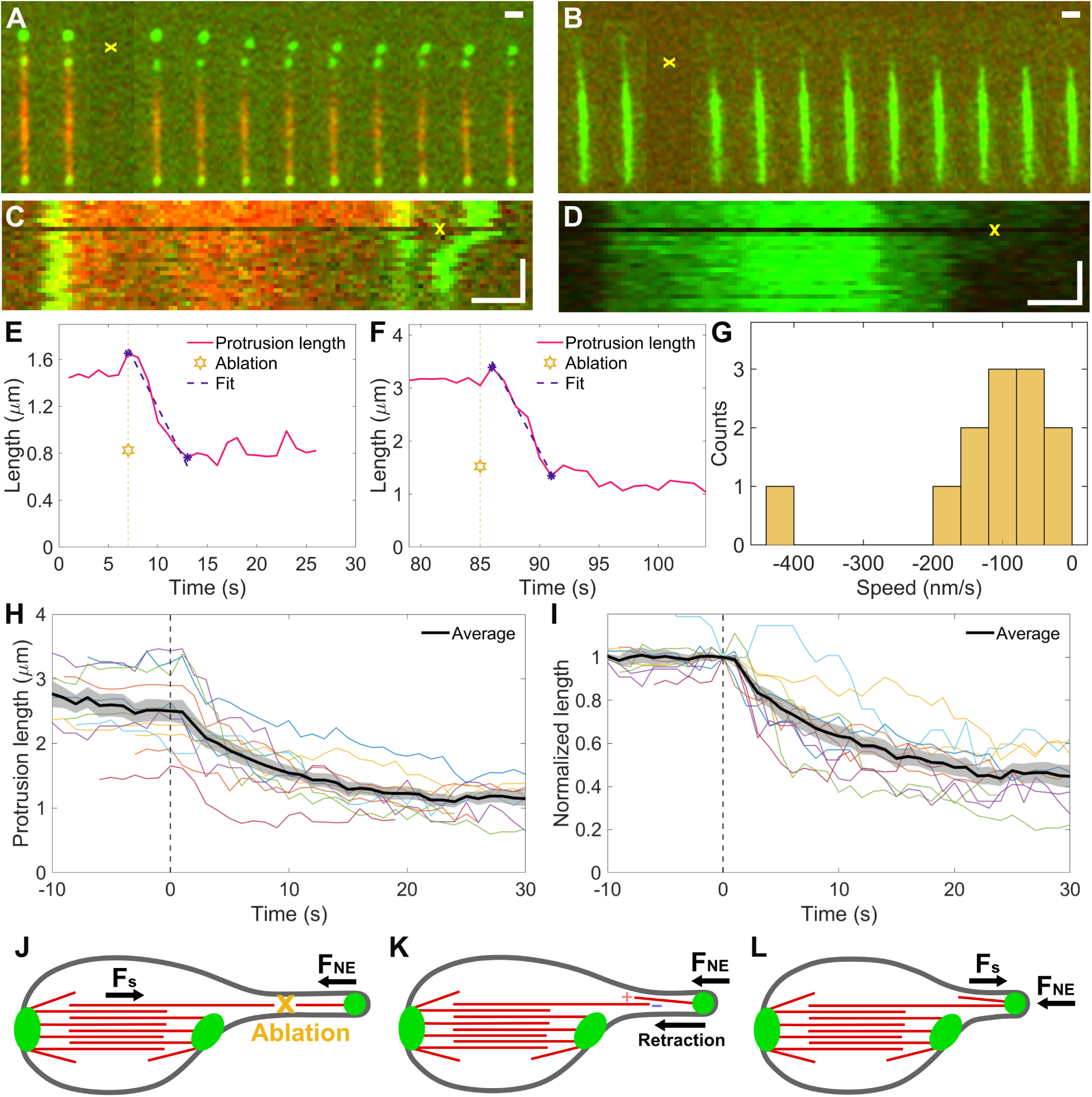
Laser ablation of protrusion microtubules causes partial protrusion retraction. **(A, B)** Fluorescence image sequences before and after laser ablation in cells showing Cut7-GFP (green) and microtubules (red, low-level mCherry-Atb2) **(A)** or microtubules (green, GFP-Atb2) **(B)**. The yellow x marks the ablation site. Time between images, 1 s; scale bar, 2 *µ*m. **(C, D)** Kymographs corresponding to the cells in A and B, where the horizontal dark line marks the time and the yellow x marks the location of laser ablation. Scale bars, 1 *µ*m and 10 s. **(E, F)** Protrusion length as a function of time for the cells shown in A and B. Gold star indicates the ablation time and location. Purple dashed line shows linear fit. **(G)** Histogram of fitted protrusion velocities after laser ablation (N = 12). **(H)** Protrusion length dynamics before and after laser ablation (*t* = 0) for *N* = 12 cells (thin colored lines), with the mean SEM shown in black and shaded gray. **(I)** Normalized protrusion length dynamics for *N* = 12 cells, with each trace normalized to 1 at *t* = 0. Black line and shaded gray are the mean SEM. **(J–L)** Conceptual models of post-ablation retraction dynamics. **(J)** Initial force balance prior to ablation (yellow x), with outward sliding force (*F*_s_) balancing inward NE force (*F*_NE_). **(K)** After ablation, the protrusion fragment experiences only *F*_NE_. Plus (+) and minus () signs indicate newly exposed microtubule ends. **(L)** Retraction stops when the protrusion fragment tip reaches the remaining microtubule.

After laser ablation near the midpoint of the protrusion, the Cut7-GFP spot at the protrusion tip retracted toward the SPB (Figure 6A and Video 5). However, this retraction typically stopped before reaching the SPB. In cells expressing GFP-Atb2, the green fluorescence signal dropped near the ablation site after laser exposure. This loss can be indicative of both photobleaching and ablation, but the subsequent partial retraction verified the protrusion microtubules were severed, not only bleached (Figure 6B).

To visualize how protrusions respond to laser ablation, we constructed kymographs. These contain a horizontal line indicating the frame during which ablation occurred (Figure 6C, D). In the Cut7-GFP kymograph, the bright protrusion tip visibly moved inward, showing retraction toward the middle SPB (Figure 6C). In the GFP-Atb2 kymograph a dim region within the protrusion microtubules, which may include both some bleached microtubules as well as the ablated region, retracted after ablation (Figure 6D).

We quantified protrusion length over time (Figure 6E, F). Individual traces showed partial retraction, with the protrusion length decreasing until the undamaged protrusion microtubules reached the site of ablation (Figure S5A–L). Linear fits to the retraction after ablation gave the retraction velocity of each event (Figure 6G), with average *−*93 *±* 13 nm/s (excluding one outlier with speed*>* 400 nm/s). This speed is not statistically significantly different from the fast retraction speed *v*_r_ *≈* 140 nm/s (Figure S3J).

To compare the distance of retraction after ablation, we aligned curves from multiple cells, setting the ablation time as *t* = 0. On average, after ablation the protrusion length decreased from 2.5 µm to 1.2 µm within 20 seconds (Figure 6H). Because initial protrusion lengths varied across cells, we normalized each trace to 1 at *t* = 0 to determine the average relative retraction distance (Figure 6I). Protrusion length remained stable during the 10 seconds prior to ablation, then rapidly decreased and plateaued around 0.5. Thus, over multiple cells protrusion microtubules partially retracted after ablation, stopping near the cut site. Less frequently, protrusions showed delayed retraction, or full retraction (Figure S5M–O).

To explain these dynamics, we envision a mechanism in which laser ablation cuts the protruding microtubules in half (Figure 6J). We expect that ablation generates a new, relatively stable microtubule minus end linked to the spindle and a less stable plus end on the remaining protrusion fragment (Figure 6K). Ablation disrupts the force balance between outward sliding and inward NE force. The protrusion fragment thus moves inward due to the NE force, until the protrusion tip contacts the newly exposed minus end (Figure 6L). This could occur simultaneously with microtubule depolymerization. We note that fission yeast spindle microtubule shortening speeds can be comparable to the retraction speeds we observed^56,58^. We analyzed the approximately one third of attempted ablations that gave a visible response of the protrusion. Other ablation events may not have cut the protrusion microtubules, or may have damaged both the microtubules and the nuclear envelope, changing the force balance.

## Discussion

Force within the mitotic spindle has been inferred in cells from the response to mechanical perturbation, including microneedle manipulation ^24,61^, laser ablation^25,48^, and spindle buckling ^13,62^, and from the stretch of structures with known mechanics such as pericentric chromatin ^63^. Spindle protrusions in fission yeast offer a complementary approach. A protrusion is a microtubule bundle of simple geometry, pushed outward by midzone motors and opposed by the force of NE deformation. Protrusions grow and shrink without external manipulation.

Comparison of our data with the force-balance model suggests an interpretation of the fast and slow protrusion length change events (Figure 7). Initial fast protrusion growth occurs when the overlap sliding force is large and not balanced by opposing force from the nuclear envelope, which could occur just after a protrusion microtubule detaches from the SPB (Figure 7A). As the protrusion extends, the overlap length drops as the NE deforms, allowing the sliding and NE forces to reach mechanical balance (Figure 7B). Subsequently, slow protrusion length changes are driven by microtubule polymerization dynamics near force balance ^42^ (Figure 7C): plus-end polymerization slightly increases the overlap length and sliding force, leading to a slight increase in protrusion length, while the opposite occurs for plus-end depolymerization. Fast retraction occurs when the sliding force is zero or greatly reduced, so that the nuclear envelope force is unbalanced (Figure 7D), which could occur if the overlap is lost or the protrusion microtubule breaks.

**Figure 7.**
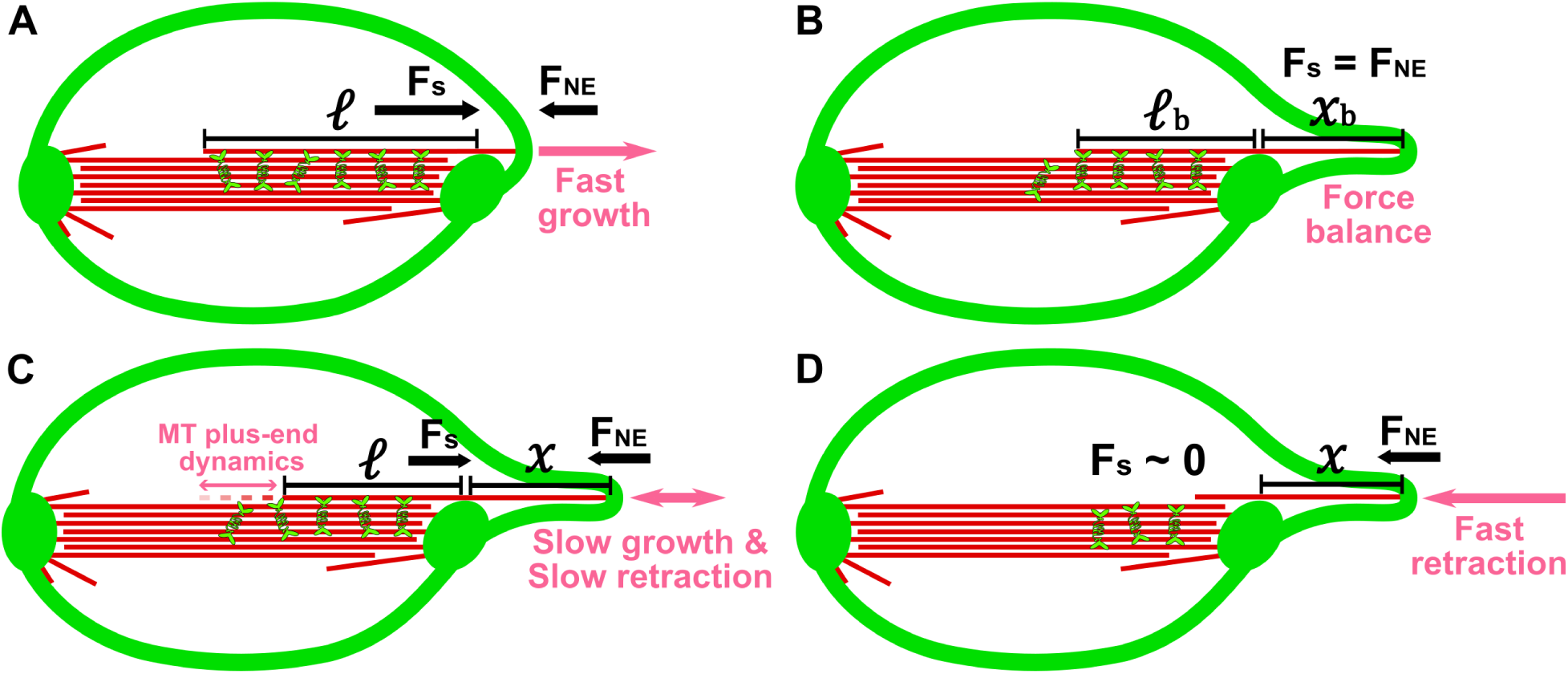
Schematic of the spindle minus-end protrusion force-balance model. **(A)** Initial fast protrusion growth, dominated by the microtubule sliding force when protrusion length is approximately zero. **(B)** Protrusion at force balance, where the sliding force equals the nuclear envelope resistance (*F*_s_ = *F*_NE_). **(C)** Subsequent slow growth and slow retraction, driven by changes in microtubule overlap length due to MT plus-end dynamics. **(D)** Fast protrusion retraction when the sliding force vanishes (*F*_s_ *≈* 0).

The kinetics of fast growth and retraction let us infer the force-balance overlap length and the protrusion microtubule length in our model, independent of NE mechanics (table 1). All inferred microtubule lengths were shorter than the spindle at protrusion onset, a check on the model. The inferred midzone sliding force density and protrusion drag coefficient depend on the NE force via the NE bending modulus. Because this modulus has not been directly measured for the fission yeast NE, we can only estimate forces to within an order of magnitude. Despite this uncertainty, the inferred midzone sliding force density is larger than that measured *in vitro* for kinesin-5 sliding apart an antiparallel microtubule pair^10,60^, even at the low end of our range. The fission yeast midzone has several antiparallel microtubule pairs^26,62^, and motors could crosslink a protrusion microtubule to more than one of them. The force per motor and the density of Cut7 in spindle overlaps have not been measured. We also note that the inferred protrusion drag coefficient is much larger than our estimate of the hydrodynamic drag on a microtubule moving through the nucleoplasm or within an NE tube, if we assume the nucleoplasm viscosity is about a factor of 500 larger than the viscosity of water. This suggests that most of the resistance to protrusion microtubule motion comes from movement of the NE and nucleus through the cytoplasm, or from friction with neighboring spindle microtubules ^64,65^.

We hypothesize that fast retraction occurs when the protrusion microtubule loses its overlap with the midzone, so that the force of NE deformation alone drives inward sliding. Microtubule depolymerization could contribute by precipitating the loss of force balance. The protrusion tip is a microtubule minus end ^38,39^, and minus ends of fission yeast spindle microtubules do not grow or shrink ^49,66,67^. The tip can therefore move toward the SPB only by sliding. Catastrophe of a plus end in the midzone, either of the protrusion microtubule or of its antiparallel partners, would shorten the overlap and reduce the sliding force. This could initiate fast retraction, with the retraction speed set by the NE force and drag. Laser ablation creates the same condition. The severed distal fragment has no overlap with the midzone, so the model predicts that it retracts at the fast retraction speed. Consistent with this expectation, the speed of retraction after laser ablation was comparable to the fast retraction speed. Because hydrodynamic drag on the microtubule is a small part of the total drag, the shorter length of the fragment should have little effect on this speed.

We found that spindle length often decreased as protrusions grew and recovered as they retracted. The amount of shortening correlated with the loss of Cut7 from the midzone. This suggests that Cut7 moving to the protrusion tip, consistent with its minus-end-directed motility^40,42,50,51,68^, reduces outward sliding within the spindle. The force of NE deformation on the protrusion could also contribute. The protrusion is coupled to the spindle through its overlap with midzone microtubules, so the spindle bears at least part of this force. Consistent with this is our observation that protrusions appear to drive spindle reorientation in some cases. Protrusions could also shorten the spindle by drawing on a limited pool of tubulin in the nucleus. Total spindle microtubule length remains constant during fission yeast mitosis, at 30 to 50 µm ^62,69^. A protrusion of one to three microtubules, each 1 to 4 µm long ^38,42^, would contain 1 to 12 µm of polymer, 2 to 40% of the spindle total. Therefore, while some protrusions may deplete the spindle tubulin pool, a typical length protrusion would deplete too little tubulin to account for the spindle shortening we observed. These three mechanisms are not mutually exclusive, and our data do not distinguish among them beyond the correlation with midzone Cut7.

Protrusions were also associated with delayed anaphase onset. Cells lacking Pkl1 but without visible protrusions entered anaphase with timing similar to wild type, so Pkl1 loss alone does not explain the delay. The delay occurred before anaphase A, while the time from anaphase A to anaphase B was unchanged. Several mechanisms could produce this delay. First, protrusions could generate erroneous kinetochore attachments that keep the spindle assembly checkpoint active. Cells lacking Pkl1 missegregate chromosomes ^31,37,39,70^. Second, spindle shortening during protrusion growth could reduce tension between sister kinetochores, which destabilizes attachments and delays biorientation ^19,71,72^. The NE force would then act indirectly, through spindle length. Third, loss of Cut7 from the midzone could slow the spindle’s progression to anaphase, independent of the NE force. Fourth, protrusions and the delay could share a common cause. Cells with more severe defects in minus-end anchoring might both form protrusions and assemble spindles more slowly.

In closed mitosis, the spindle assembles and elongates inside an intact NE, with the SPBs inserted in it. Abnormal spindles can deform the NE and cause NE division to fail^28^, while increased NE tension bends spindles, slows their elongation, and changes the protein composition of the midzone ^13,73,74^. Protrusions show that the NE also resists deformation by a single microtubule bundle, and that the spindle may respond to the force of this deformation. The asymmetric NE shapes around protrusions and the accompanying spindle rotation suggest that this force need not act along the spindle axis. Anchoring of microtubule minus ends at the SPB by the Pkl1-Msd1-Wdr8 complex ^37,38,75^ therefore has a mechanical role: it keeps outward sliding force within the spindle. Without anchoring, part of this force deforms the NE. Protrusions both reflect and perturb spindle force balance, making them a probe of the mechanical coupling between the spindle and NE in closed mitosis.

## Methods

### Cell preparation

*Schizosaccharomyces pombe* (fission yeast) strains used in this study are shown in table S1. Cell stocks were maintained using standard techniques on YE5S (Yeast Extract plus 5 Supplements) nutritionally rich media ^76^. Cells were maintained at 25 °C for standard growth and microscopy.

### Reagents and techniques

We purchased oligonucleotide primers from Integrated DNA Technologies (Coralville, IA) and restriction enzymes and Phusion HF DNA polymerase from New England Biolabs (Ipswich, MA). DNA was prepared using Qiaprep Spin Miniprep Kit and polymerase chain reaction (PCR) products were purified using Qiaquick PCR Purification Kit, both from Qiagen (Germantown, MD). Fission yeast genomic DNA was prepared using the YeaStar Genomic DNA Kit from Zymo Research (Irvine, CA). DNA sequencing was performed by Quintarabio (Hayward, CA). We determined DNA concentration using a Thermo Scientific Nanodrop 2000. Strains were verified by PCR and sequence analysis of genomic DNA compared to the *S. pombe* sequence ^77^. Cells were cultured using standard techniques ^76^.

### Strain construction

All strains were constructed via genetic crosses in the *pkl1+* or *pkl1*Δ background to study the effects of protrusions. The desired genotypes from crosses were isolated using random spore analysis and subsequent replica plating. Replica plates used YE5S plus agar with the relevant antibiotic added or Edinburgh Minimal Medium plus agar with the relevant auxotrophic marker removed from the supplemented media. The mis12-RFP marker for MB1447 was originally obtained through crossing with strain FY20781 (NBRP, Japan). The GFP-atb2 marker for MB1398 was originally obtained through crossing with the MWE2 strain from Mary Elting at North Carolina State University.

### Live-cell imaging

All microscopy images and related datasets were prepared as smoothed maximum-intensity projections using the Smooth and Z-project functions in FIJI/ImageJ. Images were obtained using live-cell preparation and time-lapse spinning-disk confocal microscopy on a Nikon Eclipse Ti microscope (Nikon USA, Melville, NY) with a Nikon 100x 1.45NA Plan Apo *λ* OIL WD 0.13 (mm) objective. Nikon NIS Elements software settings included an EM gain of 300 with exposure time of 100 ms for both the 488 nm and the 561 nm lasers. Lasers were set to 100% laser power, and 7 or 9 Z planes were acquired with 500 nm distance between each plane. The triggered acquisition setting was used to prevent photobleaching when switching between Z planes and between lasers. Z plane changes used a Physik Instruments (Auburn, MA) P-736 piezoelectric actuator. Unless otherwise noted, imaging was performed with a time of 9 seconds between each successive Z stack.

To acquire high-temporal resolution images (Figure 5), we used an Evident Scientific IXPlore IX85 SpinSR (Evident, Needham, MA) spinning-disk confocal microscope. Images were obtained using live-cell preparation and time-lapse imaging with an Evident 60X 1.50NA UPLAPO OHR Oil objective with a correction collar. The Evident cellSens software settings were exposure times of 100 ms for the 561 nm laser and 50 ms for the 488 nm laser. Lasers were set to 40% laser power for 561 nm laser and 20% for the 488 nm laser, and 5 Z planes were acquired with 500 nm distance between each plane. Z plane changes used a Märzhäuser Wetzlar piezo Z-stage (model 00-55-600-0000, Märzhäuser Wetzlar, Germany). Each image stack was obtained every 1 second for 5 minutes of total imaging.

To prepare for microscopy cells were grown on YE5S at 25 °C as previously described^19,40,41^. Cells were incubated on YE5S plates for 48 h and restreaked every 12 h. A small volume of exponentially growing cells was removed from the petri dish and placed in 10 µL of Edinburgh minimal medium (MPBio, Solon, OH) which was previously filtered with a 0.2 µm cellulose acetate filter (VWR International, Radnor, PA) to reduce background fluorescence. Cells in Edinburgh minimal medium were placed on 22×60-mm coverslips pretreated with 8 µL of lectins from *Bandeiraea simplicifolia* (Sigma-Aldrich, Burlington, MA). These coverslips were pre-equilibrated to 25 °C. For cells imaged on the Nikon microscope, temperature was maintained with *±*0.1 °C precision using a CherryTemp temperature controller (Cherry Biotech, Rennes, France) equipped with a 100X objective temperature sensor which sensed the objective temperature and incrementally adjusted the coverslip to compensate for temperature changes at the objective. For cells imaged on the Evident microscope, temperature was maintained at 25 °C in 35 mm dishes with a Tokai-HIT STX (Bala Cynwyd, PA) with a high capacity, warmed external bottle.

### Spindle and protrusion length tracking

Because more than 90% of mitotic spindles in our dataset remained straight, we measured spindle length using the distance between the two SPBs. We tracked both SPBs to subpixel accuracy using FYSKA^53^. To determine the tip position, we manually identified the protrusion’s location in each frame where protrusions were present. We then fitted a 2D Gaussian to the GFP intensity distribution to determine the protrusion GFP spot’s location. We calculated protrusion length as the distance between the middle SPB and the protrusion spot.

To measure pre-anaphase spindle length and anaphase spindle elongation speed, we manually identified the time of approximately steady pre-anaphase spindle length, the time of anaphase B onset, and the times at which steady spindle elongation began and ended, as described previously^42^. Pre-anaphase spindle length was measured using all data points within the approximately steady pre-anaphase window. Anaphase spindle elongation speed was determined from a linear fit to the region of steady elongation. To determine average spindle length as a function of time, we aligned data from each cell to anaphase B onset (Figure S1A–D). We then averaged spindle length measurements within overlapping 60-second time windows, sampled at 30-second intervals.

To define the spindle shortening distance (Figure 4I), we identified the local maximum and minimum of spindle length pre-anaphase. This was done by manually clicking a time point on the curve and searching for local extrema within five time points of the selected point. The difference between the local maximum and local minimum was the spindle shortening distance. These same time points were also used to measure spindle intensity differences (Figure S1E, F).

To determine protrusion speeds, we manually identified the start and end points of each fit (indicated by asterisks) and performed a linear least-squares fit on time points between them. Each fit included at least five data points (Figure 5C, D) and (Figure S3A–D).

### Kymograph creation and intensity analysis

For frames without protrusions, we generated two-color kymographs using FYSKA^53^ by linearly interpolating fluorescence intensity in both channels along the straight line of interest (LOI) defined by the two SPBs. When protrusions were present, we modified the LOI from a single straight line into a two-segment axis anchored at the middle SPB, while keeping the total combined length of segment one and segment two constant across all frames. To fully capture any intensity signals at and beyond both ends, we extended the LOI by a fixed length (10 pixels) at each end, beyond the far SPB and the protrusion. If the LOI were extended to the outside of the cell segmentation boundary, interpolating across these areas where no fluorescence data existed resulted in zero-intensity regions on the kymograph (Video 1). A rectangular box was drawn along the LOI to enclose both spindle poles and the spindle midzone as the spindle region, and another box was drawn along the LOI to enclose the protrusion and its connection to the nearest spindle pole as the protrusion region (Video 2). Pixel intensity in each region was summed to find the total spindle intensity and total protrusion intensity.

### Quantification of anaphase onset time

To quantify the time of anaphase onset, we imaged strains expressing Cut7-GFP, Mis12-RFP, and Sid4-mCherry in the *pkl1+* and *pkl1* Δ backgrounds (table S1). We acquired images every 20 s for 45 min. We analyzed cells for which mitotic entry and anaphase onset were observed within this time. We manually identified mitotic entry, defined as the first appearance of Cut7-GFP fluorescence at the SPBs; anaphase A onset, identified by the start of movement of Mis12-RFP signal to the two SPBs; and anaphase B onset, defined as the beginning of rapid SPB separation (Figure 3A). Because the Mis12-RFP signals were sometimes too dim to reliably identify kinetochore separation, cells with ambiguous anaphase A onset were excluded from analysis of anaphase A timing. For *pkl1* Δ cells we classified cells as protrusion (P) and no-protrusion (NP), where cells were denoted P if a protrusion was visible at any time between mitotic entry and anaphase onset. The time from mitotic entry to anaphase A onset, from mitotic entry to anaphase B onset, and from anaphase A onset to anaphase B onset were quantified and compared among *pkl1+ pkl1* Δ (NP), and *pkl1* Δ (P) cells using one-way ANOVA followed by Fisher’s least significant difference (LSD) post-hoc tests.

### Protrusion length dynamics model and fitting to data

A force-balance model was found to capture protrusion dynamics in previous work ^42^. In this model, the total length of a protrusion microtubule is *L* = *x* + *ℓ*, where *x* is the length of the part of the protrusion that extends beyond the SPB, and *ℓ* is the length of the segment that overlaps the spindle midzone. We assume that midzone motors push the protrusion outward (increasing *x* and decreasing *ℓ*) with sliding force *F*_s_, while the NE tube pushes the protrusion inward with force *F*_NE_. We also assume that the protrusion microtubule has a drag coefficient *γ*. Then the protrusion equation of motion is

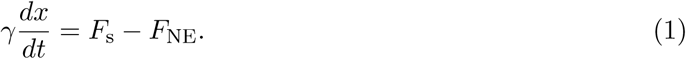

Based on the observation that protrusion length can change at both fast and slow speeds, we assume that fast length changes occur when the net force is nonzero and the protrusion is not in mechanical equilibrium. During these rapid changes, we assume that the protrusion microtubule length *L* is constant. Protrusion microtubules undergo dynamic instability that contributes to length changes ^42^, but polymerization occurs relatively slowly on the time scale we are considering. Therefore, during fast events we assume that any changes in *x* occur because sliding trades off length between the overlap and the protrusion. When *L* is fixed, the rate of change of *x* reaches zero when the forces balance at mechanical equilibrium, at the force-balance values we denote *x*_b_ and *ℓ*_b_.

To determine when force balance is reached, we assume consistent with previous work ^42^ that midzone motors produce a net force proportional to overlap length *F*_s_ = *f*_s_*ℓ,* where *f*_s_ is the sliding force per unit length. Then, with a constant NE force, force balance occurs when *f*_s_*ℓ*_b_ = *F*_NE_ and *L* = *x*_b_ + *ℓ*_b_. Using these results and *ℓ* = *L − x*, we can rewrite the protrusion length dynamics equation as

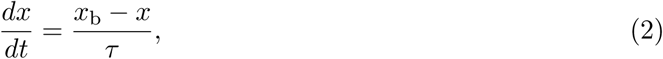

where 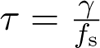. If the initial condition is *x*(*t* = *t*_0_) = *x*_0_, the solution is

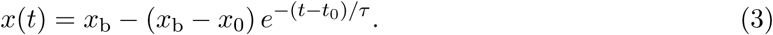

Note that this exponential solution depends on the assumption that *γ* is constant, independent of *x* and *ℓ*. We discuss the limits of this assumption below.

### Nuclear envelope force, bending modulus, and surface tension

The NE consists of two lipid bilayers fused at nuclear pores and separated by a lumen about 32 nm wide ^59^. We model the NE as a single surface with effective bending modulus *κ* and tension *σ*, each summed over the two bilayers. The corresponding single bilayer values are *κ*_1_ and *σ*_1_. For a tubular NE extension longer than its radius (*x > R*), the NE force is constant^19,42,78,79^,

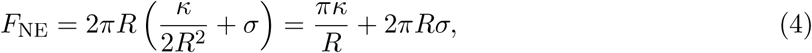

with bending modulus *κ*, surface tension *σ*, and tube radius *R*. The full expression ^78^ is within 1% of this value for *x >* 190 nm at *R* = *R*_0_. The preferred radius of an NE tube *R_0_ = √κ/2σ*^80^.

For the fission yeast NE, only the ratio of tension to bending modulus has been measured. The value *σ/κ* = 65 µm*^−^*^2^ was determined by matching computed shapes to electron micrographs of an interphase nucleus deformed by an intranuclear microtubule bundle ^59^. To obtain the bending modulus and surface tension, prior work estimated a typical value of the bending modulus *κ* = 4.00 10*^−^*^19^ J = 400 pN nm, which gives *σ* = 0.026 pN*/*nm for the whole NE and *σ*_1_ = 0.013 pN*/*nm per bilayer ^59^. The per-bilayer tension is the same as the tension of ER membranes measured by tether pulling ^81^.

We estimate a range of possible values for *κ* from measurements on single bilayers and modeling. For vesicles measured by micropipette aspiration, the bending modulus ranges from 40 pN nm to 120 pN nm ^82,83^. Measurements from vesicle shape fluctuations are higher than micropipette measurements ^83,84^. Most fluctuation measurements fall between 100 and 160 pN nm, while the full range is 25 to 246 pN nm ^83,84^. For biological membranes, tether pulling from ER and Golgi membranes gave *κ*_1_ = 330 pN nm ^81^. The spacing between the two membranes may make the NE stiffer than two independent bilayers. Separated membranes support larger bending moments, and in modeling two membranes 45 nm apart and fused at pores 350 nm apart had an effective bending modulus of 1700 pN nm, 10 times that of one membrane ^85^. This stiffening depends on geometry, and ranges from 2 to 10 times the single-membrane value ^85^. This calculation was for a point force on a flat patch, not a tube as in our NE deformations.

Our previous work on spindle buckling may constrain the upper end of this range ^13^. In unperturbed cells spindles rarely buckle, while in cerulenin-treated cells spindles buckle at lengths near 5 µm, where the buckling force is about 31 pN ^13,62^. If we assume that the force scale for deforming the NE *f_0_ = 2π√2κσ* stays below the buckling force in unperturbed cells, then using the measured ratio we find *f_0_ = 2πκσ2σ/κ = (0.0716nm^-1^)κ*, so that *κ <* 430 pN nm. Note that this bound is uncertain, because of uncertainty in the buckling prefactor, the single measurement of *σ/κ*, and the shape of the NE near the spindle poles.

Given this uncertainty, we take *κ* to range from 80 to 660 pN nm, with an estimate 230 pN nm, the geometric middle of the range, and the corresponding *σ* to range from 0.005 to 0.043 pN/nm, with an estimate 0.015 pN/nm. We chose the upper end somewhat above the buckling bound to allow for its uncertainty. This value corresponds to two ER-like membranes without stiffening from the membrane spacing. With our estimates *κ* = 230 pN nm, *σ* = 0.015 pN/nm, and *R* = *R*_0_ = 88 nm, we find *F*_NE_ = 16.5 pN. This is an order-of-magnitude estimate: the range of bending moduli we are considering could increase or decrease this force by a factor of 3.

### Drag coefficient

We hypothesize that fast retraction occurs when the protrusion microtubule loses contact with the midzone, so *F*_s_ = 0. The retraction speed is then (eqn. (1)) 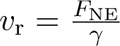. The measured mean speed was *v*_r_ = 140 nm*/*s, so *γ* = *F*_NE_*/v*_r_ *≈* 118 pN s*/*µm, with range 41 to 340 pN s/µm depending on the value of *κ*.

To determine whether a constant *γ* assumption is justified, we consider three likely sources of drag. First, we consider drag of a microtubule parallel to its axis. For a rod of length *L* and diameter *d* = 25 nm in a fluid of viscosity *η*, 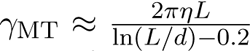. If *L* is fixed, *γ*_MT_ is constant during a fast retraction event. Next, we consider drag of a microtubule in the NE tube. For a single microtubule of radius *a* = 12.5 nm sliding inside a tube of radius *R*_0_ = 88 nm, shear in the fluid between the microtubule and tube gives a drag proportional to 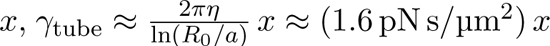 *x ≈* (1.6 pN s*/*µm^2^) *x* assuming the nucleoplasm viscosity is about a factor of 500 larger than the viscosity of water^42^. This term changes with *x* during an event, but by about 0.8 pN s/µm for a change in *x* of 0.5 µm, less than 1% of the drag coefficient we estimate. Third, drag could arise from crosslinkers and motors in the overlap that resist sliding. Previous work found crosslinker friction *in vitro* that depends on crosslinker number, not overlap length^65^. This term is absent during fast retraction, so the *γ* estimated from *v*_r_ does not include it.

The microtubule terms together are under 4 pN s/µm, about 3% of our estimate of *γ* and under 10% at the low end of the range, and their variation within an event is a few percent. The remaining *≈* 115 pN s*/*µm (range 38 to 335 pN s/µm) likely comes from the NE (motion of the tube and nucleus through the cytoplasm, or membrane flow into the tube) or from friction with neighboring spindle microtubules ^64,65^.

### Fit results and derived forces

To compare the model to fast protrusion length change events, we fit eqn. (3) to selected growth events to determine *x*_b_ and *τ*. We fit only initial fast growth events, which began at protrusion length below 0.5 µm, lasted less than 10 s, and showed a clear transition to slow growth. Of the 35 fast growth events, 14 were not initial events and were excluded. We fit 9 of the remaining 21 events, excluding those with less pronounced fast growth. We performed the fits with the scipy.optimize.least squares function using the Trust Region Reflective algorithm. Growth events often began with a brief period when protrusion length dipped or flattened, which the model does not describe. We excluded this period from the fits. It may reflect interactions with the SPB, which are not included in the model. Then given the retraction speed *v*_r_, we find the force-balance overlap length *ℓ*_b_ = *v*_r_*τ*, the protrusion microtubule length *L* = *x*_b_ +*ℓ*_b_, and the sliding force density 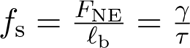. Of these inferred quantities, only *γ* and *f*_s_ depend on *F*_NE_.

The fits gave mean *τ* = 2.4 s and *x*_b_ = 1.0 µm. With *v*_r_ = 140 nm*/*s, the mean force-balance overlap length is *ℓ*_b_ = 0.33 µm and the mean protrusion microtubule length is *L* = 1.4 µm. With *F*_NE_ = 16.5 pN, the sliding force density is *f*_s_ *≈* 50 pN*/*µm, with range 17 to 140 pN/µm (Table 1).

### Laser ablation of protruding microtubules

We performed spinning disk confocal live imaging and laser ablation experiments as described previously^13^. We captured time-lapse images using a Nikon Ti-E stand on an Andor Dragon-fly spinning disk confocal fluorescence microscope containing a spinning disk dichroic Chroma ZT405/488/561/640rpc, 488 nm (50 mW) diode laser (240 ms exposures) with Borealis attachment (Andor), emission filter Chroma ET525/50m, and an Andor iXon3 camera. We performed imaging with a 100x 1.45 Ph3 Nikon objective and a 1.5x magnifier (built-in to the Dragonfly system).

We used an Andor Micropoint attachment with galvo-controlled steering for targeted laser ablation, delivering 20-30 ns pulses of 551 nm light at 20 Hz. We imaged each cell until either protrusion reformation and continued elongation had been completed or the protrusion had failed to repair after 5 min. For protrusion ablation videos, we imaged every 1 s. We used Andor Fusion software to manage imaging and Andor IQ software to simultaneously manipulate the laser ablation system.

Following laser ablation imaging, we manually labeled the locations of SPBs and protrusion tips frame-by-frame due to low signal-to-noise ratios (S/N) or the absence of Sid4-marked SPBs. After manual annotation of these spot positions, we used FYSKA to generate kymographs^53^. For individual cells, we plotted protrusion length as a function of time, aligning each individual curve to the time of ablation (*t* = 0) to construct the ensemble average. Additionally, we normalized each individual curve of protrusion length as a function of time to 1 at *t* = 0 to calculate the mean normalized curve (Figure 6I).

### Online supplemental material

Figure S1 shows spindle length over time in individual cells and additional examples of pre-anaphase spindle shortening. Figure S2 shows additional examples of protrusion dynamics. Figure S3 shows linear fits of protrusion length changes, protrusion velocity distributions, and retraction speeds with and without laser ablation. Figure S4 shows fits of the force-balance model to individual fast growth events. Figure S5 shows protrusion length after laser ablation for individual cells and atypical responses to ablation. Table S1 lists the strains used. Videos 1 and 2 show the line of interest and the intensity boxes used for kymograph analysis. Videos 3 and 4 show a kymograph being constructed from the Cut7-GFP and Sid4-mCherry channels of one cell. Video 5 shows protrusion retraction after laser ablation.

## Data availability

Data are available at the CU Scholar repository at (DOI to be added). Simulation code is available on Github at (link to be added).

## Acknowledgements

We thank Andrew Berger, Wilson Lough, and Brittany Wu for helpful discussions, Christian Pagan Medina for help with cell collection and sample preparation for laser ablation experiments, and James Orth for assistance with microscopy. This work is funded by grants NIH R01GM164613 and NSF MCB 2133243 to MDB, and NIH R35GM138083 and NSF MCB 2133276 to MWE. We acknowledge the Light Microscopy Core Facility in Porter Biosciences at the University of Colorado Boulder (RRID:SCR 018993) for help and advice with microscopy. Spinning disk microscopy was supported by NIH shared equipment grant S10OD038323.

## Disclosures

The authors declare no competing interests exist.

## Author contributions

B. Zhou: Data curation, Formal analysis, Investigation, Methodology, Resources, Software, Validation, Visualization, Writing - original draft, Writing - review & editing, Z.R. Gergely: Data curation, Formal analysis, Investigation, Methodology, Resources, Supervision, Validation, Visualization, Writing - original draft, D.G. Steckhahn: Conceptualization, Data curation, Formal analysis, Investigation, Methodology, Software, Validation, Visualization, Writing - original draft, T. Mahoney: Investigation, Resources, Writing - original draft, L. Swallow: Formal analysis, Investigation, J.R. McIntosh: Methodology, Resources, Supervision, Visualization, Writing - original draft, M.W. Elting: Funding acquisition, Investigation, Methodology, Project administration, Resources, Supervision, Validation, Visualization, Writing - original draft, M.D. Betterton: Conceptualization, Formal analysis, Funding acquisition, Methodology, Project administration, Resources, Supervision, Visualization, Writing - original draft, Writing - review & editing.

---

**Figure S1.**
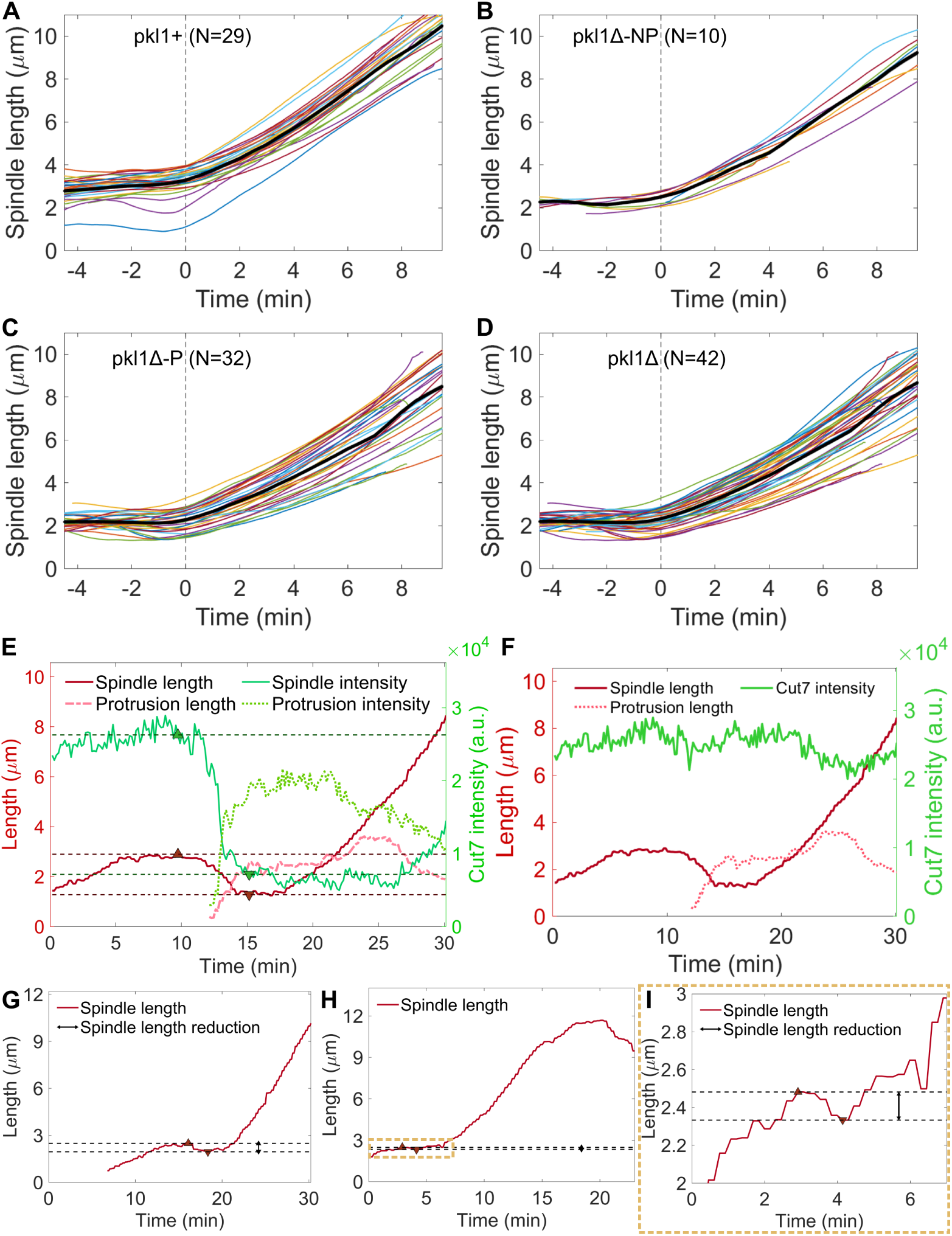
Spindle length as a function of time in individual cells and additional examples showing spindle length shortening during pre-anaphase. (A–D) Spindle length of individual cells (thin colored lines) for *pkl1+* (*N* = 29) **(A)**, *pkl1* Δ-NP (*N* = 10) **(B)**, *pkl1* Δ-P (*N* = 32) **(C),** and all *pkl1* Δ (*N* = 42) **(D)** cells, aligned to anaphase B onset (*t* = 0, vertical dashed line). Black lines are the mean, and gray shading is the SEM. **(E)** Spindle and protrusion length (left axis, red) and Cut7-GFP intensity on the spindle and protrusion (right axis, green) as a function of time. Upward and downward dark-red triangles indicate local maxima and minima. Dashed lines indicate the spindle shortening distance and change in Cut7-GFP intensity. **(F)** Spindle and protrusion length (left axis, red) and total Cut7-GFP intensity (right axis, green) as a function of time for the cell shown in E. **(G, H)** Pre-anaphase spindle length reduction in two example *pkl1* Δ cells without a visible protrusion. The golden dashed box in H indicates the region enlarged in I. **(I)** Enlarged view of the pre-anaphase spindle length reduction for the cell shown in H.

**Figure S2.**
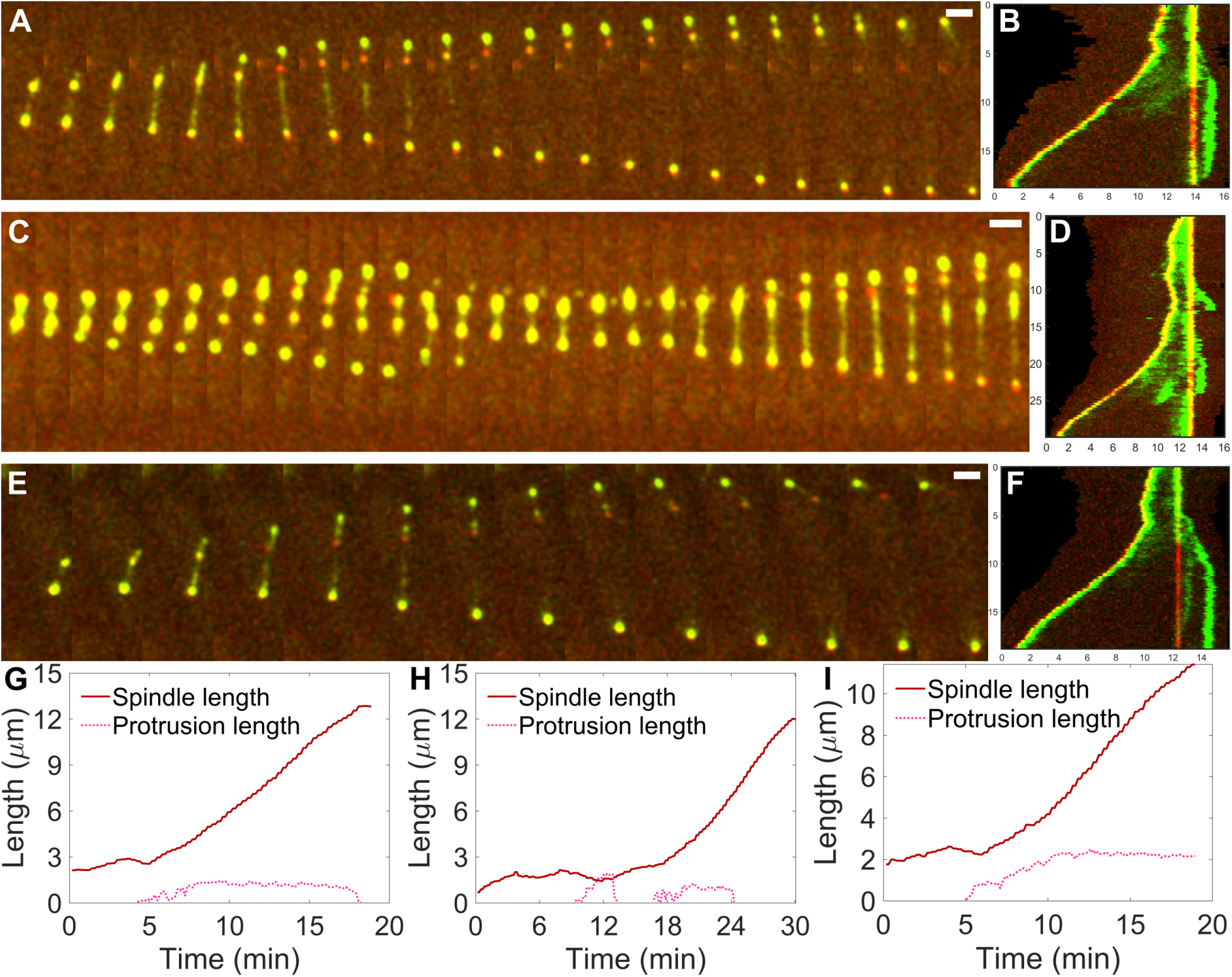
Spindle microtubule minus-end protrusion dynamics. **(A, C, E)** Image sequences of *pkl1* Δ cells showing protrusion dynamics. **(B, D, F)** Two-color kymographs of the cells shown in A, C, and E, showing SPBs (red, Sid4-mCherry) and Cut7-GFP (green). **(G–I)** Spindle and protrusion length as a function of time for the cells shown in A and B **(G)**, C and D **(H)**, and E and F **(I)**. Time between images, 38 s in A, 39 s in C, and 76 s in E; scale bars, 2 *µ*m.

**Figure S3.**
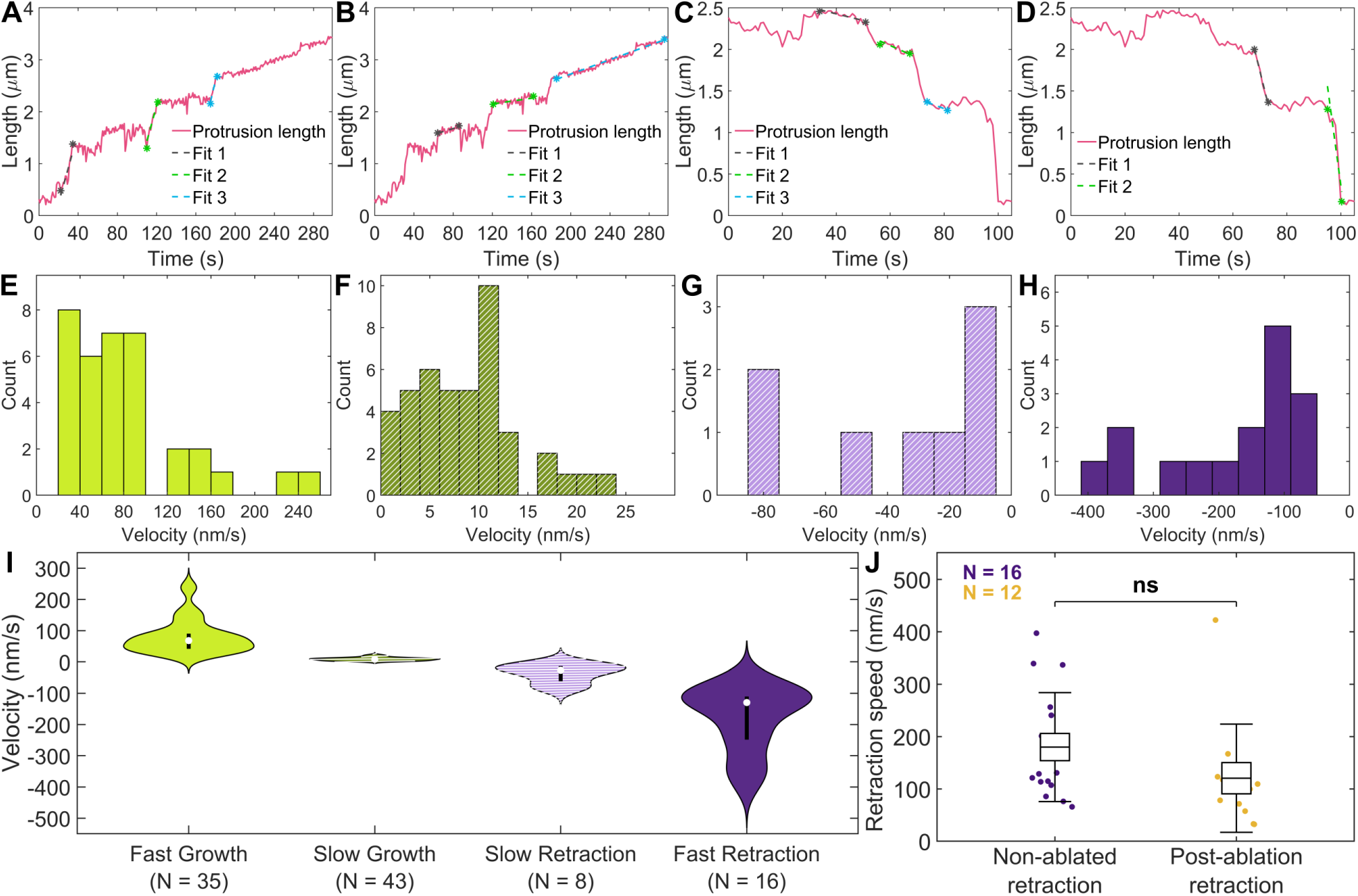
Piecewise linear fits to determine protrusion growth and retraction speeds and protrusion velocity distributions. (A–D) Protrusion length as a function of time for two *pkl1* Δ cells, showing fast (A) and slow (B) protrusion growth, and slow (C) and fast (D) retraction. Asterisks mark the start and end of each fitted interval. **(E–H)** Histograms of protrusion length change events showing fast growth (E, bin size: 20 nm/s), slow growth (F, bin size: 2 nm/s), slow retraction (G, bin size: 10 nm/s), and fast retraction (H, bin size: 40 nm/s), as in fig. 5E. **(I)** Violin plot of the velocity distributions. **(J)** Comparison of protrusion retraction speed without laser ablation (*N* = 16) and following laser ablation (*N* = 12). Boxes show the mean and SEM; error bars show the standard deviation. Statistical significance was determined using one-way ANOVA; ns = not significant.

**Figure S4.**
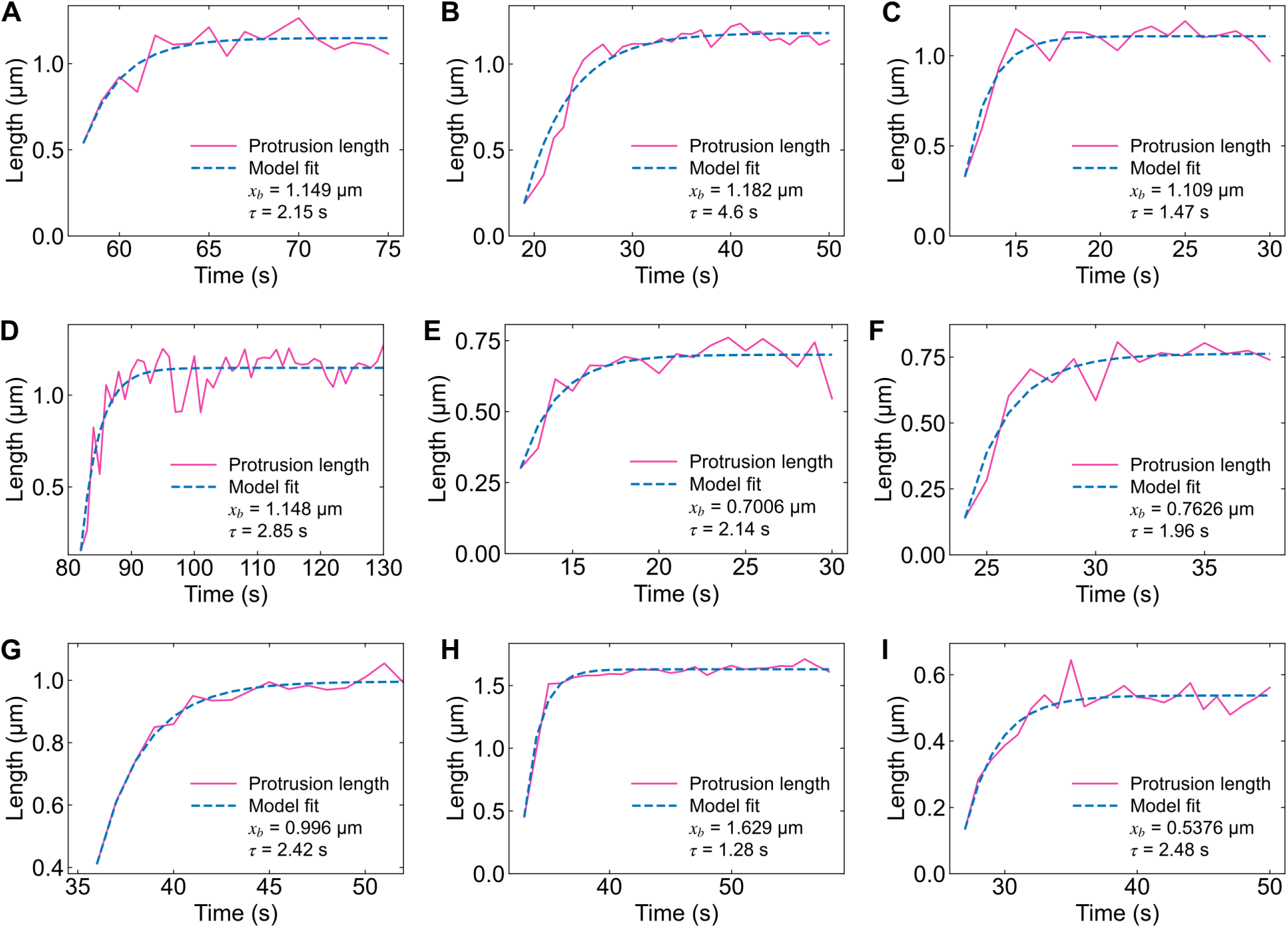
Fits of the force-balance model to fast protrusion growth. (A–I) Individual protrusion growth traces (pink solid lines) and force-balance model fits (blue dashed lines), showing the two fit parameters force balance protrusion length (*x*_b_) and timescale (*τ*).

**Figure S5.**
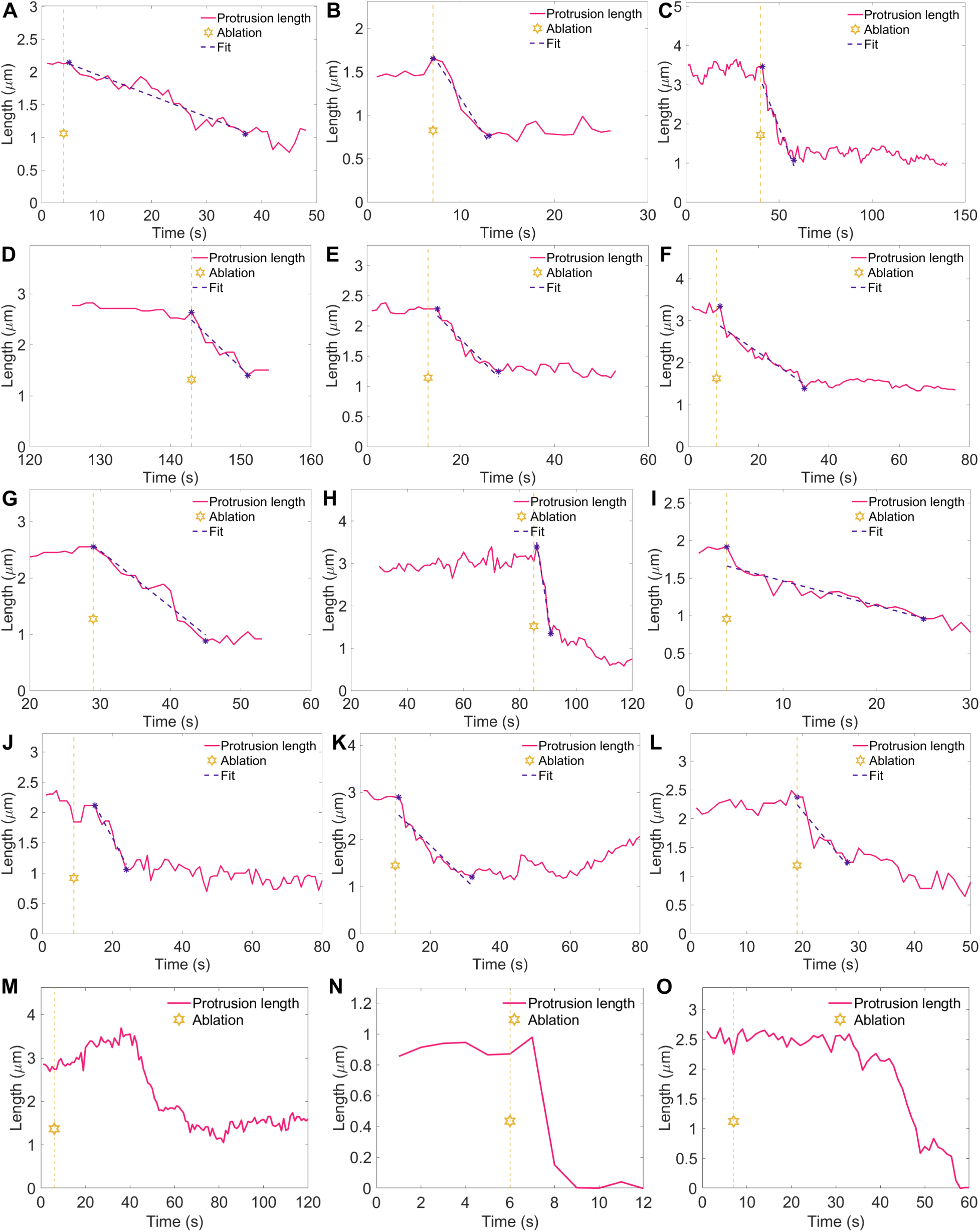
Laser ablation of protrusion microtubules, fitting of partial protrusion retraction speeds, and atypical protrusion responses following laser ablation. (A–L) Protrusion length as a function of time for cells (*N* = 12) showing partial protrusion retraction. Gold stars indicate the time of laser ablation, and purple dashed lines show linear fits. **(M–O)** Atypical protrusion responses to laser ablation. (M) Delayed partial retraction (*N* = 4). **(N)** Full retraction (*N* = 1). **(O)** Delayed full retraction (*N* = 2).

**Table S1.** List of strains, their genotypes, and associated figures.

| Strain (MB) | Genotype | Figures |
|---|---|---|
| 1316 | cut7-GFP-KanR, sid4-mCh:hphMX6, his3<sup>−</sup>, leu1-32, ura4<sup>−</sup>, ade6? | Figure 1, S1 |
| 1302 | cut7-GFP-KanR, sid4-mCh:hphMX6, pkl1Δ::his3<sup>+</sup>, leu1-32, his3<sup>−</sup>, ura4-D18, ade6? | Figure 1, 4, 5, S1–S4 |
| 1409 | cut11-GFP:ura4+, z:adh15:mCherry-atb2:natMX6, pkl1D::his3+, his3-, leu1-32, ura4-D18, ade6? | Figure 2 |
| 1445 | mis12-RFP::LEU2, cut7-GFP-kanR, sid4-mCh:hphMX6, pkl1D::his3+, his3-, leu1-32, ura4-D18, ade6? | Figure 3 |
| 1447 | mis12-RFP::LEU2, cut7-GFP-kanR, sid4-mCh:hphMX6, pkl1+, his3-, leu1-32, ura4-D18, ade6? | Figure 3 |
| 1150 | pkl1D::his3+, cut7-GFP-kanr, z:adh15:mCherry-atb2:natMX6 pku80::hphMX6, leu?, his- | Figure 6, S3, S5 |
| 1151 | pkl1D::his3+, cut7-GFP-kanr, z:adh15:mCherry-atb2:natMX6, leu?, his- | Figure 6, S3, S5 |
| 1398 | GFP-atb2:kanMX6, sid4-mCh:hphMX6, pkl1D::his3+ leu1-32, his3-D1, ura4-D18, ade6? | Figure 6, S3, S5 |

## Video legends

**Video 1.** Two-segment line of interest used to construct kymographs, shown on the merged channels of a *pkl1* Δ cell. Frames collected every 9.5 s. Video playback is 10 frames per second.

**Video 2.** Boxes used to measure spindle and protrusion Cut7-GFP intensity, for the cell in Video 1. Frames collected every 9.5 s. Video playback is 10 frames per second.

**Video 3.** Kymograph construction from the Cut7-GFP channel of a *pkl1* Δ cell. Frames collected every 9.5 s. Video playback is 10 frames per second.

**Video 4.** Kymograph construction from the Sid4-mCherry channel of the cell in Video 3. Frames collected every 9.5 s. Video playback is 10 frames per second.

**Video 5.** Protrusion retraction after laser ablation in a cell expressing Cut7-GFP and mCherry-Atb2. Frames collected every 1 s. Video playback is 10 frames per second.

## Notes

### Competing Interest Statement

The authors have declared no competing interest.

## References

[1] Gul Civelekoglu-Scholey and Jonathan M. Scholey. Mitotic force generators and chromosome segregation. Cellular and Molecular Life Sciences, 67(13):2231–2250, July 2010.

[2] S. Dumont and T. Mitchison. Mechanical forces in mitosis. Comprehensive Biophysics, 4, 2011.

[3] J. Richard McIntosh. Mitosis. Cold Spring Harbor Perspectives in Biology, 8(9):a023218, September 2016.

[4] Nenad Pavin and Iva M. Tolić. Self-Organization and Forces in the Mitotic Spindle. Annual Review of Biophysics, 45(1):279–298, 2016.

[5] Mary Williard Elting, Pooja Suresh, and Sophie Dumont. The Spindle: Integrating Architecture and Mechanics across Scales. Trends in Cell Biology, 28(11):896–910, November 2018.

[6] David J. Sharp, Kent L. McDonald, Heather M. Brown, Heinrich J. Matthies, Claire Walczak, Ron D. Vale, Timothy J. Mitchison, and Jonathan M. Scholey. The Bipolar Kinesin, KLP61F, Cross-links Microtubules within Interpolar Microtubule Bundles of Drosophila Embryonic Mitotic Spindles. The Journal of Cell Biology, 144(1):125–138, November 1999.

7. Lukas C. Kapitein, Erwin J. G. Peterman, Benjamin H. Kwok, Jeffrey H. Kim, Tarun M. Kapoor, and Christoph F. Schmidt. The bipolar mitotic kinesin Eg5 moves on both microtubules that it crosslinks. Nature, 435(7038):114–118, May 2005.

[8] A. Mogilner and E. Craig. Towards a quantitative understanding of mitotic spindle assembly and mechanics. Journal of Cell Science, 123(20):3435–3445, 2010.

[9] Viktoriya Syrovatkina, Chuanhai Fu, and Phong T. Tran. Antagonistic Spindle Motors and MAPs Regulate Metaphase Spindle Length and Chromosome Segregation. Current Biology, 23(23):2423–2429, December 2013.

[10] Yuta Shimamoto, Scott Forth, and Tarun M. Kapoor. Measuring Pushing and Braking Forces Generated by Ensembles of Kinesin-5 Crosslinking Two Microtubules. Developmental Cell, 34(6):669–681, September 2015.

[11] Robert Blackwell, Christopher Edelmaier, Oliver Sweezy-Schindler, Adam Lamson, Zachary R. Gergely, Eileen O’Toole, Ammon Crapo, Loren E. Hough, J. Richard McIntosh, Matthew A. Glaser, and Meredith D. Betterton. Physical determinants of bipolar mitotic spindle assembly and stability in fission yeast. Science Advances, 3(1):e1601603, January 2017.

[12] Benjamin Lacroix and Julien Dumont. Spatial and Temporal Scaling of Microtubules and Mitotic Spindles. Cells, 11(2):248, January 2022.

[13] Marcus A. Begley, Taylor Mahoney, Christian Pagán Medina, Parsa Zareiesfandabadi, Matthew B. Rapp, Mastawal Tirfe, Sharonda J. LeBlanc, Meredith D. Betterton, and Mary Williard Elting. Mechanical Coupling With the Nuclear Envelope Shapes the Schizosaccharomyces pombe Mitotic Spindle. Cytoskeleton, n/a(n/a), 2025.

[14] Venecia A. Valdez, Lila Neahring, Sabine Petry, and Sophie Dumont. Mechanisms underlying spindle assembly and robustness. Nature Reviews Molecular Cell Biology, 24(8):523–542, August 2023.

[15] E.N. Cytrynbaum, J.M. Scholey, and A. Mogilner. A Force Balance Model of Early Spindle Pole Separation in Drosophila Embryos. Biophysical Journal, 84(2):757–769, February 2003.

[16] Li Tao, Alex Mogilner, Gul Civelekoglu-Scholey, Roy Wollman, James Evans, Henning Stahlberg, and Jonathan M. Scholey. A Homotetrameric Kinesin-5, KLP61F, Bundles Microtubules and Antagonizes Ncd in Motility Assays. Current Biology, 16(23):2293–2302, December 2006.

[17] Christian Hentrich and Thomas Surrey. Microtubule organization by the antagonistic mitotic motors kinesin-5 and kinesin-14. The Journal of Cell Biology, 189(3):465–480, March 2010.

[18] Masashi Yukawa, Yusuke Yamada, Tomoaki Yamauchi, and Takashi Toda. Two spatially distinct Kinesin-14 Pkl1 and Klp2 generate collaborative inward forces against Kinesin-5 Cut7 in S. pombe. J Cell Sci, 131:jcs.210740, 2018.

[19] Christopher Edelmaier, Adam R Lamson, Zachary R Gergely, Saad Ansari, Robert Blackwell, J Richard McIntosh, Matthew A Glaser, and Meredith D Betterton. Mechanisms of chromosome biorientation and bipolar spindle assembly analyzed by computational modeling. eLife, 9:e48787, February 2020.

[20] Masashi Yukawa, Yasuhiro Teratani, and Takashi Toda. How Essential Kinesin-5 Becomes Non-Essential in Fission Yeast: Force Balance and Microtubule Dynamics Matter. Cells, 9(5):1154, May 2020.

[21] Parsa Zareiesfandabadi and Mary Williard Elting. Force by minus-end motors Dhc1 and Klp2 collapses the S. pombe spindle after laser ablation. Biophysical Journal, 121(2):263–276, January 2022.

[22] Gohta Goshima, Shigeaki Saitoh, and Mitsuhiro Yanagida. Proper metaphase spindle length is determined by centromere proteins Mis12 and Mis6 required for faithful chromosome segregation. Genes & Development, 13(13):1664–1677, January 1999.

[23] R. B. Nicklas. Measurements of the force produced by the mitotic spindle in anaphase. The Journal of Cell Biology, 97(2):542–548, January 1983.

[24] Yuta Shimamoto, Yusuke T. Maeda, Shin’ichi Ishiwata, Albert J. Libchaber, and Tarun M. Kapoor. Insights into the Micromechanical Properties of the Metaphase Spindle. Cell, 145(7):1062–1074, June 2011.

[25] Mary Williard Elting, Christina L. Hueschen, Dylan B. Udy, and Sophie Dumont. Force on spindle microtubule minus ends moves chromosomes. Journal of Cell Biology, 206(2):245–256, July 2014.

[26] R. Ding, K. L. McDonald, and J. R. McIntosh. Three-dimensional reconstruction and analysis of mitotic spindles from the yeast, Schizosaccharomyces pombe. The Journal of Cell Biology, 120(1):141–151, January 1993.

[27] I. M. Hagan. The fission yeast microtubule cytoskeleton. Journal of Cell Science, 111(12):1603–1612, June 1998.

[28] Liling Zheng, Cindi Schwartz, Valentin Magidson, Alexey Khodjakov, and Snezhana Oliferenko. The spindle pole bodies facilitate nuclear envelope division during closed mitosis in fission yeast. PLoS biology, 5(7):e170, July 2007.

[29] Iain Hagan and Mitsuhiro Yanagida. Novel potential mitotic motor protein encoded by the fission yeast cut7+ gene. Nature, 347(6293):563–566, October 1990.

[30] Chuanhai Fu, Jonathan J. Ward, Isabelle Loiodice, Guilhem Velve-Casquillas, Francois J. Nedelec, and Phong T. Tran. Phospho-Regulated Interaction between Kinesin-6 Klp9p and Microtubule Bundler Ase1p Promotes Spindle Elongation. Developmental Cell, 17(2):257–267, August 2009.

[31] A L Pidoux, M LeDizet, and W Z Cande. Fission yeast pkl1 is a kinesin-related protein involved in mitotic spindle function. Molecular Biology of the Cell, 7(10):1639–1655, October 1996.

[32] Cynthia L. Troxell, Mark A. Sweezy, Robert R. West, Karen D. Reed, Bryan D. Carson, Alison L. Pidoux, W. Zacheus Cande, and J. Richard McIntosh. Pkl1 +andklp2 +: Two Kinesins of the Kar3 Subfamily in Fission Yeast Perform Different Functions in Both Mitosis and Meiosis. Molecular Biology of the Cell, 12(11):3476–3488, January 2001.

[33] Sergio A. Rincon, Adam Lamson, Robert Blackwell, Viktoriya Syrovatkina, Vincent Fraisier, Anne Paoletti, Meredith D. Betterton, and Phong T. Tran. Kinesin-5-independent mitotic spindle assembly requires the antiparallel microtubule crosslinker Ase1 in fission yeast. Nature Communications, 8:15286, May 2017.

[34] Masashi Yukawa, Masaki Okazaki, Yasuhiro Teratani, Ken’ya Furuta, and Takashi Toda. Kinesin-6 Klp9 plays motor-dependent and -independent roles in collaboration with Kinesin-5 Cut7 and the microtubule crosslinker Ase1 in fission yeast. Scientific Reports, 9(1):7336, May 2019.

[35] Zachary T. Olmsted, Timothy D. Riehlman, Carmen N. Branca, Andrew G. Colliver, Leilani O. Cruz, and Janet L. Paluh. Kinesin-14 Pkl1 targets *γ*-tubulin for release from the *γ*-tubulin ring complex (*γ*-TuRC). Cell Cycle, 12(5):842–848, March 2013.

[36] Zachary T. Olmsted, Andrew G. Colliver, Timothy D. Riehlman, and Janet L. Paluh. Kinesin-14 and kinesin-5 antagonistically regulate microtubule nucleation by *γ*-TuRC in yeast and human cells. Nature Communications, 5:5339, October 2014.

[37] Masashi Yukawa, Chiho Ikebe, and Takashi Toda. The Msd1–Wdr8–Pkl1 complex anchors microtubule minus ends to fission yeast spindle pole bodies. J Cell Biol, 209(4):549–562, May 2015.

[38] Mika Toya, Masamitsu Sato, Uta Haselmann, Kazuhide Asakawa, Damian Brunner, Claude Antony, and Takashi Toda. *γ*-Tubulin complex-mediated anchoring of spindle microtubules to spindle-pole bodies requires Msd1 in fission yeast. Nature Cell Biology, 9(6):646–653, June 2007.

[39] Viktoriya Syrovatkina and Phong T. Tran. Loss of kinesin-14 results in aneuploidy via kinesin-5-dependent microtubule protrusions leading to chromosome cut. Nature Communications, 6:7322, June 2015.

[40] Zachary R. Gergely, Saad Ansari, Michele H. Jones, Bojun Zhou, Cai Cash, Richard McIntosh, and Meredith D. Betterton. Kinesin-5/Cut7 moves bidirectionally on fission-yeast spindles with activity that increases in anaphase. Journal of Cell Science, 136:jcs.260474, January 2023.

[41] Zachary R. Gergely, Michele H. Jones, Bojun Zhou, Cai Cash, J. Richard McIntosh, and Meredith D. Betterton. Distinct regions of the kinesin-5 C-terminal tail are essential for mitotic spindle midzone localization and sliding force. Proceedings of the National Academy of Sciences, 120(39):e2306480120, September 2023.

[42] Michele H. Jones, Zachary R. Gergely, Daniel Steckhahn, Bojun Zhou, and Meredith D. Betterton. Kinesin-5/Cut7 C-terminal tail phosphorylation is essential for microtubule sliding force and bipolar mitotic spindle assembly. Current Biology, 34(20):4781–4793.e6, October 2024.

[43] Candice Yam, Yue He, Dan Zhang, Keng-Hwee Chiam, and Snezhana Oliferenko. Divergent strategies for controlling the nuclear membrane satisfy geometric constraints during nuclear division. Current biology: CB, 21(15):1314–1319, August 2011.

[44] Stefania Castagnetti, Bojan Bozic, and Sasa Svetina. Mechanical and molecular basis for the symmetrical division of the fission yeast nuclear envelope. Physical Chemistry Chemical Physics, 17(24):15629–15636, 2015.

[45] Qian Zhu, Fan Zheng, Allen P. Liu, Jin Qian, Chuanhai Fu, and Yuan Lin. Shape Transformation of the Nuclear Envelope during Closed Mitosis. Biophysical journal, 111(10):2309–2316, November 2016.

[46] Ai Takemoto, Shigehiro A. Kawashima, Juan-Juan Li, Linda Jeffery, Kenzo Yamatsugu, Olivier Elemento, and Paul Nurse. Nuclear envelope expansion is crucial for proper chromosomal segregation during a closed mitosis. Journal of Cell Science, 129(6):1250–1259, March 2016.

[47] Chao Fang, Fan Zheng, Jiaxing Yao, Xi Wei, Chuanhai Fu, Xinghua Shi, and Yuan Lin. A model for bridging microtubule dynamics with nuclear envelope shape evolution during closed mitosis. Journal of the Mechanics and Physics of Solids, 144:104116, November 2020.

[48] Iva M Tolić-Norrelykke, Leonardo Sacconi, Geneviève Thon, and Francesco S Pavone. Positioning and Elongation of the Fission Yeast Spindle by Microtubule-Based Pushing. Current Biology, 14(13):1181–1186, July 2004.

[49] Alexey Khodjakov, Sabrina La Terra, and Fred Chang. Laser Microsurgery in Fission Yeast: Role of the Mitotic Spindle Midzone in Anaphase B. Current Biology, 14(15):1330–1340, August 2004.

[50] Masaki Edamatsu. Bidirectional motility of the fission yeast kinesin-5, Cut7. Biochemical and Biophysical Research Communications, 446(1):231–234, March 2014.

[51] Mishan Britto, Adeline Goulet, Syeda Rizvi, Ottilie von Loeffelholz, Carolyn A. Moores, and Robert A. Cross. Schizosaccharomyces pombe kinesin-5 switches direction using a steric blocking mechanism. Proceedings of the National Academy of Sciences, 113:201611581, November 2016.

[52] Lora Winters, Ivana Ban, Marcel Prelogović, Iana Kalinina, Nenad Pavin, and Iva M. Tolić. Pivoting of microtubules driven by minus-end-directed motors leads to spindle assembly. BMC Biology, 17(1):42, May 2019.

[53] Bojun Zhou, Zachary R. Gergely, Michele H. Jones, and Meredith D. Betterton. Tracking Mitotic Spindle Dynamics and Protein Localization in Fission Yeast With FYSKA, the Fission Yeast Spindle Kymograph Analyzer. *Cytoskeleton*, n/a(n/a), 2026.

[54] Austin J. Hepperla, Patrick T. Willey, Courtney E. Coombes, Breanna M. Schuster, Maryam Gerami-Nejad, Mark McClellan, Soumya Mukherjee, Janet Fox, Mark Winey, David J. Odde, Eileen O’Toole, and Melissa K. Gardner. Minus-End-Directed Kinesin-14 Motors Align Antiparallel Microtubules to Control Metaphase Spindle Length. Developmental Cell, 31(1):61–72, October 2014.

[55] Maria Makarova, Ying Gu, Jun-Song Chen, Janel Renée Beckley, Kathleen Louise Gould, and Snezhana Oliferenko. Temporal Regulation of Lipin Activity Diverged to Account for Differences in Mitotic Programs. Current Biology, 26(2):237–243, January 2016.

[56] M. J. Sagolla, S. Uzawa, and W. Z. Cande. Individual microtubule dynamics contribute to the function of mitotic and cytoplasmic arrays in fission yeast. Journal of Cell Science, 116(24):4891–4903, 2003.

[57] Iana Kalinina, Amitabha Nandi, Petrina Delivani, Mariola R. Chacón, Anna H. Klemm, Damien Ramunno-Johnson, Alexander Krull, Benjamin Lindner, Nenad Pavin, and Iva M. Tolić-Norrelykke. Pivoting of microtubules around the spindle pole accelerates kinetochore capture. Nature Cell Biology, 15:82–87, 2012.

[58] Robert Blackwell, Oliver Sweezy-Schindler, Christopher Edelmaier, Zachary R. Gergely, Patrick J. Flynn, Salvador Montes, Ammon Crapo, Alireza Doostan, J. Richard McIntosh, Matthew A. Glaser, and Meredith D. Betterton. Contributions of Microtubule Dynamic Instability and Rotational Diffusion to Kinetochore Capture. Biophysical Journal, 112(3):552–563, February 2017.

[59] Gerald H. W. Lim, Greg Huber, Yoshihiro Torii, Aiko Hirata, Jonathan Miller, and Shelley Sazer. Vesicle-Like Biomechanics Governs Important Aspects of Nuclear Geometry in Fission Yeast. PLoS ONE, 2(9):e948, September 2007.

[60] Tatyana Bodrug, Elizabeth M Wilson-Kubalek, Stanley Nithianantham, Alex F Thompson, April Alfieri, Ignas Gaska, Jennifer Major, Garrett Debs, Sayaka Inagaki, Pedro Gutierrez, Larisa Gheber, Richard J McKenney, Charles Vaughn Sindelar, Ronald Milligan, Jason Stumpff, Steven S Rosenfeld, Scott T Forth, and Jawdat Al-Bassam. The kinesin-5 tail domain directly modulates the mechanochemical cycle of the motor domain for anti-parallel microtubule sliding. eLife, 9:e51131, January 2020.

[61] Alexandra F. Long, Pooja Suresh, and Sophie Dumont. Individual kinetochore-fibers locally dissipate force to maintain robust mammalian spindle structure. Journal of Cell Biology, 219(8):e201911090, May 2020.

[62] Jonathan J. Ward, Hélio Roque, Claude Antony, and Fraņcois Nédélec. Mechanical design principles of a mitotic spindle. eLife, 3:e03398, January 2015.

[63] Andrew D. Stephens, Rachel A. Haggerty, Paula A. Vasquez, Leandra Vicci, Chloe E. Snider, Fu Shi, Cory Quammen, Christopher Mullins, Julian Haase, Russell M. Taylor, Jolien S. Verdaasdonk, Michael R. Falvo, Yuan Jin, M. Gregory Forest, and Kerry Bloom. Pericentric chromatin loops function as a nonlinear spring in mitotic force balance. The Journal of Cell Biology, 200(6):757–772, March 2013.

[64] Andrew Ward, Feodor Hilitski, Walter Schwenger, David Welch, A. W. C. Lau, Vincenzo Vitelli, L. Mahadevan, and Zvonimir Dogic. Solid friction between soft filaments. Nature Materials, 14(6):583–588, June 2015.

[65] Ignas Gaska, Mason E. Armstrong, April Alfieri, and Scott Forth. The Mitotic Crosslinking Protein PRC1 Acts Like a Mechanical Dashpot to Resist Microtubule Sliding. Developmental Cell, 54(3):367–378.e5, August 2020.

[66] Aneil Mallavarapu, Ken Sawin, and Tim Mitchison. A switch in microtubule dynamics at the onset of anaphase B in the mitotic spindle of Schizosaccharomyces pombe. Current Biology, 9(23):1423–1428, December 1999.

[67] Sabina Zimmerman and Fred Chang. Effects of *γ*-Tubulin Complex Proteins on Microtubule Nucleation and Catastrophe in Fission Yeast. Molecular Biology of the Cell, 16(6):2719–2733, January 2005.

[68] M. Edamatsu. Molecular properties of the N-terminal extension of the fission yeast kinesin-5, Cut7. Genetics and molecular research: GMR, 15(1), February 2016.

[69] Isabelle Loiodice, Marcel E. Janson, Penny Tavormina, Sebastien Schaub, Divya Bhatt, Ryan Cochran, Julie Czupryna, Chuanhai Fu, and Phong T. Tran. Quantifying Tubulin Concentration and Microtubule Number Throughout the Fission Yeast Cell Cycle. Biomolecules, 9(3), March 2019.

[70] Ekaterina L. Grishchuk, Ilia S. Spiridonov, and J. Richard McIntosh. Mitotic Chromosome Biorientation in Fission Yeast Is Enhanced by Dynein and a Minus-end–directed, Kinesin-like Protein. Molecular Biology of the Cell, 18(6):2216–2225, January 2007.

[71] Hilary Dewar, Kozo Tanaka, Kim Nasmyth, and Tomoyuki U. Tanaka. Tension between two kinetochores suffices for their bi-orientation on the mitotic spindle. Nature, 428(6978):nature02328, February 2004.

[72] Shelby L. McVey, Jenna K. Cosby, and Natalie J. Nannas. Aurora B Tension Sensing Mechanisms in the Kinetochore Ensure Accurate Chromosome Segregation. International Journal of Molecular Sciences, 22(16):8818, January 2021.

[73] Taylor Mahoney, Christopher Needham, Reem Hakeem, Marcus Begley, Christian Pagán Medina, and Mary Williard Elting. Spindle biochemistry responds to compressive force from the nuclear envelope to tune spindle dynamics during closed mitosis. Preprint, bioRxiv, January 2026.

[74] Paula Real-Calderón, Thomas G. Fai, Rafael R. Daga, Joël Lemière, and Fred Chang. Analyses of bent spindles reveal the mechanics of anaphase B in fission yeast. Molecular Biology of the Cell, pages mbc.E25–11–0556, February 2026.

[75] Priyanka Sasmal, Makito Miyazaki, Frédérique Carlier-Grynkorn, and Phong T. Tran. Chimeras of kinesin-6 and kinesin-14 reveal head-neck-tail domain functions and dysfunctions that lead to aneuploidy in fission yeast. Current Biology, 36(2):415–425.e5, January 2026.

[76] Sergio Moreno, Amar Klar, and Paul Nurse. Molecular genetic analysis of fission yeast Schizosaccharomyces pombe. In Guide to Yeast Genetics and Molecular Biology, volume Volume 194, pages 795–823. Academic Press, 1991.

[77] V. Wood, R. Gwilliam, M.-A. Rajandream, M. Lyne, R. Lyne, A. Stewart, J. Sgouros, N. Peat, J. Hayles, S. Baker, D. Basham, S. Bowman, K. Brooks, D. Brown, S. Brown, T. Chillingworth, C. Churcher, M. Collins, R. Connor, A. Cronin, P. Davis, T. Feltwell, A. Fraser, S. Gentles, A. Goble, N. Hamlin, D. Harris, J. Hidalgo, G. Hodgson, S. Holroyd, T. Hornsby, S. Howarth, E. J. Huckle, S. Hunt, K. Jagels, K. James, L. Jones, M. Jones, S. Leather, S. McDonald, J. McLean, P. Mooney, S. Moule, K. Mungall, L. Murphy, D. Niblett, C. Odell, K. Oliver, S. O’Neil, D. Pearson, M. A. Quail, E. Rabbinowitsch, K. Rutherford, S. Rutter, D. Saunders, K. Seeger, S. Sharp, J. Skelton, M. Simmonds, R. Squares, S. Squares, K. Stevens, K. Taylor, R. G. Taylor, A. Tivey, S. Walsh, T. Warren, S. Whitehead, J. Woodward, G. Volckaert, R. Aert, J. Robben, B. Grymonprez, I. Weltjens, E. Vanstreels, M. Rieger, M. Schäfer, S. Müller-Auer, C. Gabel, M. Fuchs, C. Fritzc, E. Holzer, D. Moestl, H. Hilbert, K. Borzym, I. Langer, A. Beck, H. Lehrach, R. Reinhardt, T. M. Pohl, P. Eger, W. Zimmermann, H. Wedler, R. Wambutt, B. Purnelle, A. Goffeau, E. Cadieu, S. Dréano, S. Gloux, V. Lelaure, S. Mottier, F. Galibert, S. J. Aves, Z. Xiang, C. Hunt, K. Moore, S. M. Hurst, M. Lucas, M. Rochet, C. Gaillardin, V. A. Tallada, A. Garzon, G. Thode, R. R. Daga, L. Cruzado, J. Jimenez, M. Sánchez, F. del Rey, J. Benito, A. Domínguez, J. L. Revuelta, S. Moreno, J. Armstrong, S. L. Forsburg, L. Cerrutti, T. Lowe, W. R. McCombie, I. Paulsen, J. Potashkin, G. V. Shpakovski, D. Ussery, B. G. Barrell, and P. Nurse. The genome sequence of Schizosaccharomyces pombe. Nature, 415(6874):871–880, February 2002.

[78] Imre Derényi, Frank Jülicher, and Jacques Prost. Formation and Interaction of Membrane Tubes. Physical Review Letters, 88(23):238101, May 2002.

[79] Adam R. Lamson, Christopher J. Edelmaier, Matthew A. Glaser, and Meredith D. Betterton. Theory of Cytoskeletal Reorganization during Cross-Linker-Mediated Mitotic Spindle Assembly. Biophysical Journal, 116(9):1719–1731, May 2019.

[80] Patricia Bassereau, Benoit Sorre, and Aurore Lévy. Bending lipid membranes: Experiments after W. Helfrich’s model. Advances in Colloid and Interface Science, 208:47–57, June 2014.

[81] Arpita Upadhyaya and Michael P. Sheetz. Tension in Tubulovesicular Networks of Golgi and Endoplasmic Reticulum Membranes. Biophysical Journal, 86(5):2923–2928, May 2004.

[82] W. Rawicz, K. C. Olbrich, T. McIntosh, D. Needham, and E. Evans. Effect of Chain Length and Unsaturation on Elasticity of Lipid Bilayers. Biophysical Journal, 79(1):328–339, July 2000.

[83] Derek Marsh. Elastic curvature constants of lipid monolayers and bilayers. Chemistry and Physics of Lipids, 144(2):146–159, November 2006.

[84] Rumiana Dimova. Recent developments in the field of bending rigidity measurements on membranes. Advances in Colloid and Interface Science, 208:225–234, June 2014.

[85] Ashutosh Agrawal and Tanmay P. Lele. Geometry of the nuclear envelope determines its flexural stiffness. Molecular Biology of the Cell, 31(16):1815–1821, July 2020.

